# Fitting individual-based models of spatial population dynamics to long-term monitoring data

**DOI:** 10.1101/2022.09.26.509574

**Authors:** Anne-Kathleen Malchow, Guillermo Fandos, Urs G. Kormann, Martin U. Grüebler, Marc Kéry, Florian Hartig, Damaris Zurell

## Abstract

Generating spatial predictions of species distribution is a central task for research and policy. Currently, correlative species distribution models (cSDMs) are among the most widely used tools for this purpose. However, cSDMs fundamental assumption of species distributions in equilibrium with their environment is rarely met in real data and limits their applicability for dynamic projections. Process-based, dynamic SDMs (dSDMs) promise to overcome these limitations as they explicitly represent transient dynamics and enhance spatio-temporal transferability. Software tools for implementing dSDMs become increasingly available, yet their parameter estimation can be complex.

Here, we test the feasibility of calibrating and validating a dSDM using long-term monitoring data of Swiss red kites (*Milvus milvus*). This population has shown strong increases in abundance and a progressive range expansion over the last decades, indicating a non-equilibrium situation. We construct an individual-based model with the RangeShiftR modelling platform and use Bayesian inference for model calibration. This allows the integration of heterogeneous data sources, such as parameter estimates from published literature as well as observational data from monitoring schemes, with coherent assessment of parameter uncertainty. Our monitoring data encompass counts of breeding pairs at 267 sites across Switzerland over 22 years. We validate our model using a spatial-block cross-validation scheme and assess predictive performance with a rank-correlation coefficient.

Our model showed very good predictive accuracy of spatial projections and represented well the observed population dynamics over the last two decades. Results suggest that reproductive success was a key factor driving the observed range expansion. According to our model, the Swiss red kite population fills large parts of its current range but has potential for further increases in density.

We demonstrate the practicality of data integration and validation for dSDMs using RangeShifteR. This approach can improve predictive performance compared to cSDMs. The workflow presented here can be adopted for any population for which some prior knowledge on demographic and dispersal parameters as well as spatio-temporal observations of abundance or presence/absence are available. The fitted model provides improved quantitative insights into the ecology of a species, which may greatly help conservation and management actions.

**Open Research statement:** This submission uses novel code which is provided in an external repository. All data and code required to replicate the presented analyses are provided as private-for-peer review via a public GitHub repository under the following link: https://github.com/UP-macroecology/Malchow_IBMcalibration_2023

Upon acceptance, this repository will be archived and versioned with Zenodo and its DOI provided.

For this study, a tagged development version of the R package RangeShiftR was used that is available at: https://github.com/RangeShifter/RangeShiftR-package/releases/tag/v.1.1-beta.0

## 1 Introduction

In response to multiple anthropogenic pressures and environmental shifts, the abundance and distribution of many species are changing (Selwood et al., 2015; Newbold et al., 2015; Díaz et al., 2019). Decreasing populations can potentially be stabilised or may even recover by effective conservation measures (Hoffmann et al., 2010; Bolam et al., 2021; Duarte et al., 2020). But also expanding populations can be the focus of conservation interest, for example when exploring scenarios of future threats or evaluating the invasive potential of a species (Thompson et al., 2021). The basis for efficient conservation planning thus lies in reliable knowledge about the spatio-temporal patterns of abundance and the expected effects of conservation measures (Guisan et al., 2013; Zurell et al., 2022).

Various approaches have been developed for spatially-explicit population modelling, ranging from purely correlative to detailed mechanistic species distribution models (SDMs; Dormann et al., 2012; Guisan et al., 2013). Currently, most spatial model assessments for conservation planning are based on projections of correlative SDMs (cSDMs) (Franklin, 2013; Zurell et al., 2022), which statistically relate species occurrences to environmental predictors (Elith & Leathwick, 2009). This class of models can achieve high flexibility and may be readily fitted to available occurrence data, but their geographical and temporal transferability is limited (Araújo & Peterson, 2012; Wenger & Olden, 2012). Furthermore, they only provide stationary or time-implicit projections, which rely on the assumption that the observed distribution stays in equilibrium with its environment, even if the environmental conditions change (Guisan & Thuiller, 2005). However, this assumption is commonly violated in conservation-relevant cases such as for invasive species, in reintroduction programs, or for threatened populations affected by ongoing environmental change. This leads to inaccurate predictions because the true species distribution is actually transient and thus dependent on time and history (Santos et al., 2020; Watts et al., 2020; Semper-Pascual et al., 2021).

Such dynamic abundance patterns can be represented with spatially-explicit process-based SDMs (hereafter called dSDMs). These models explain present species distributions and abundances as the result of interacting ecological processes, such as demography, dispersal and evolution (Urban et al., 2016). To this end, dSDMs explicitly describe at least one of these processes to model spatio-temporal and potentially transient population dynamics. Examples include representations of local population dynamics (Keith et al., 2008; Barber-O’Malley et al., 2022) and limiting processes like dispersal (Risk et al., 2011; Broms et al., 2016; Smolik et al., 2010; Hefley et al., 2017; Wikle, 2003), physiology (Rodríguez et al., 2019), or species interactions (Schweiger et al., 2012; Pellissier et al., 2013). Further, dSDMs often include stochastic elements to account for processes not explicitly described by the model or for intrinsic variability. Thanks to the integration of ecological theory, dSDMs are expected to provide more accurate predictions under extrapolation and thus to be more readily transferable to non-analog conditions than cSDMs (Gallien et al., 2010).

A challenge for working with dSDMs is their specification and validation (Schmolke et al., 2010). Fully specifying a dSDM includes two main steps, both of which require distinct types of knowledge about the population of interest (Singer et al., 2018; Fig. 1). First, in the model building step, the model structure and the functional description of the relevant processes are established. Both are usually chosen based on ecological theory and expert opinion. Second, in the parameterisation step, the numeric values of all model parameters, such as demographic and dispersal rates, are determined by direct or inverse (also called indirect) parameterisation or a combination of both. Direct parameterisation uses estimates of process parameters based on data collected in the field or from experiments. Inverse parameterisation determines likely parameter values by comparing the generated model response, typically spatio-temporal abundance or occurrence, with observed field data. To make efficient use of all sources of information and combine both direct and inverse parameterisation, a Bayesian calibration framework can be employed. In this, the direct parameterisation and its uncertainty are expressed as prior distributions. The prior is updated via Bayes’ rule using a likelihood which measures how well a given set of parameter values is able to reproduce the observed response data. This updated prior yields the posterior distribution (Hobbs & Hooten, 2015). The procedure thus identifies parameterisations that are consistent with the data and can generate new knowledge on the studied population, as prior estimates of process parameters are modified by new data and their uncertainty may be reduced. This approach further allows the consistent propagation of uncertainty from the data sources through to model projections (Hartig et al., 2012; Marion et al., 2012; Jaatinen et al., 2021). Ultimately, in the validation step, the predictive performance of the specified model is assessed. For this, the model is evaluated on a set of testing data that, ideally, is independent of the training data. One way to generate the training and testing data is cross-validation, in which the full data set is partitioned in a prescribed way, e.g. in leave-p-out or k-fold cross-validation (Arlot & Celisse, 2010). The final, validated model can be used to generate projections to other times or places and to compare the outcome of alternative management scenarios (Bleyhl et al., 2021).

**Figure 1:**
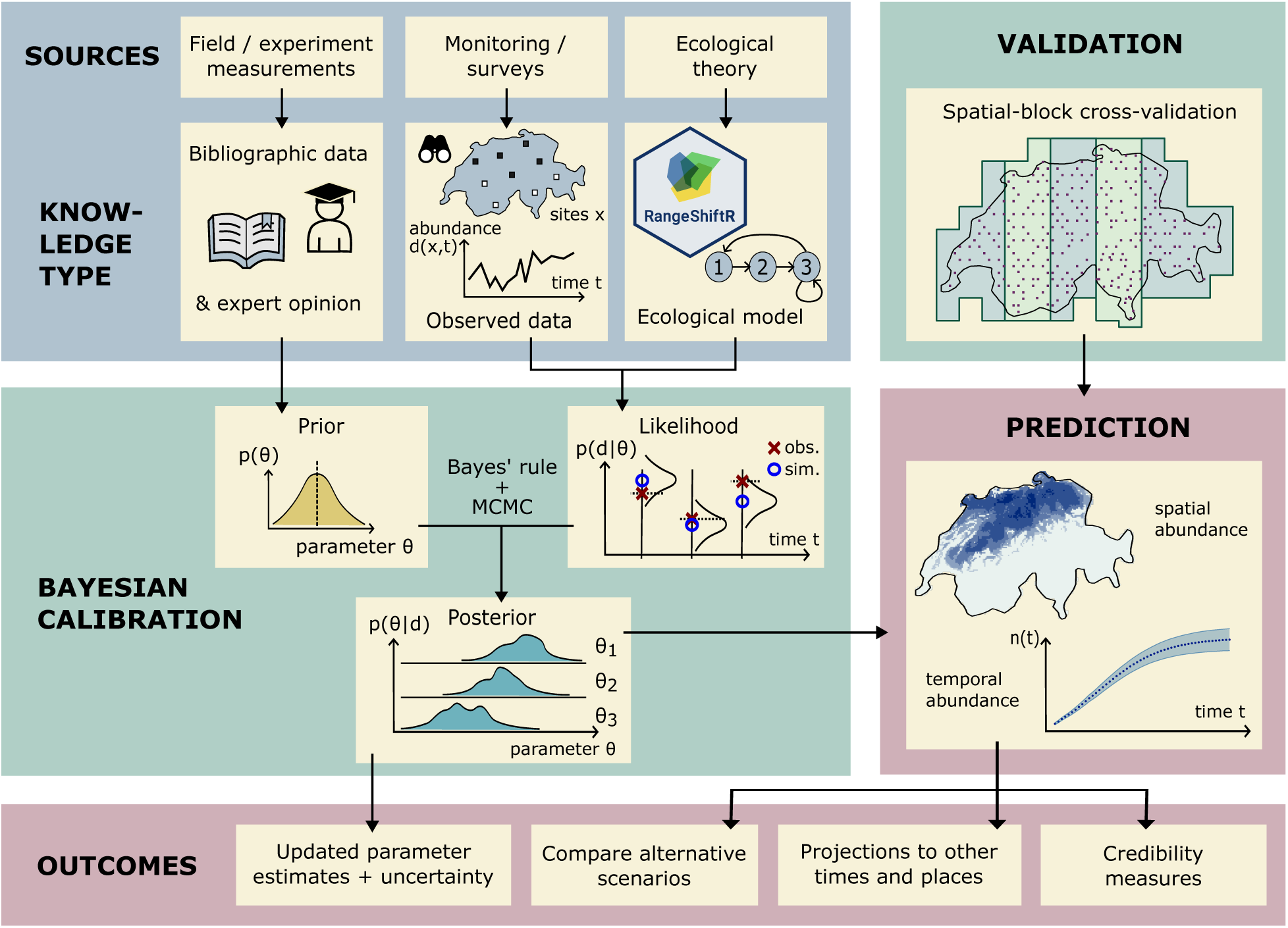
Calibration and validation workflow for process-based, dynamic distribution and population models. Different types of knowledge are needed to specify the model structure and parameterisation. Their direct specification can be informed from literature data, expert opinion, and ecological theory. In a calibration, the direct knowledge on model parameters is combined with observations of the modelled response. When using Bayesian inference, this is done via the likelihood function evaluated for this data. For cross-validation, the calibration is repeated for different subsets of data, using the held-out data to measure predictive performance. Multiple outcomes can be derived, both from the posterior distribution directly and from model projections.

Despite repeated calls for more dynamic SDM approaches, the widespread use of dSDMs for conservation applications has been hampered by technical challenges with respect to their parameterisation and validation (Briscoe et al., 2019). With the proliferation of novel methods for the various model building steps, software tools are being developed that assist their case-specific implementation. In R, these are available as packages for building different types of complex dSDMs (Visintin et al., 2020; Malchow et al., 2021; Fordham et al., 2021; Moulin et al., 2021; Landguth et al., 2017; Hagen et al., 2021), for model calibration (Hartig et al., 2019; Csilléry et al., 2012), and for cross-validation (Valavi et al., 2019). However, their combined application in integrated modelling workflows is still challenging and rarely done.

In this study, we present a complete calibration and validation workflow for dSDMs, utilising heterogeneous data for direct and inverse parameterisation. As a case study, we modelled the Swiss population of red kite (*Milvus milvus*). This population has a highly dynamic recent history with accelerating increases over the last 50 years (Aebischer & Scherler, 2021), rendering a dynamic modelling approach adequate. We first built a dSDM with the individual-based modelling (IBM) platform RangeShiftR which explicitly simulates the processes of population dynamics and dispersal (Malchow et al., 2021). Then, its process parameters were directly parameterised using published literature data (Table S1). This direct parameterisation was subsequently updated by integrating information from long-term, structured survey data with Bayesian inference using the R package BayesianTools (Hartig et al., 2019). Finally, the predictive performance of the calibrated model was evaluated by cross-validation on spatially-blocked data folds (D. R. Roberts et al., 2017). To test our workflow, we investigated whether the calibration could successfully inform parameter estimates and which process parameters were most sensitive to the survey data. Comparing the prior and posterior predictive distributions under our model, we assessed whether the calibration improved model fit and reduced uncertainty. Predictive performance of the fitted model was evaluated in spatial-block cross-validation and compared to a cSDM fitted to the same data. Lastly, the calibrated model was used to explore the potential population size and distribution of red kite in Switzerland.

Our workflow (Fig. 1) is intended to guide the application of complex dSDMs to populations that exhibit substantial dynamics and for which suitable data sources for direct and inverse parameterisation are available. It is suited for linking process-based models with monitoring data to obtain a solid quantitative basis for management decisions while being explicit about all uncertainties involved (Zylstra & Zipkin, 2021).

## 2 Materials and Methods

### 2.1 Data

We utilised two sources of monitoring data of the red kite in Switzerland: the Swiss breeding bird atlas that provides snapshot data from two periods (1993-1996, Schmid et al., 1998; and 2013-2016, Knaus et al., 2018) and the Swiss breeding bird survey (MHB, Schmid et al., 2004) that provides abundance time series for every year since 1999. We used the MHB series up to the year 2019. Both schemes are based on so-called simplified territory mapping of a systematic-random sample 1 km^2^ squares across Switzerland and record the number of observed breeding pairs during two to three repeat surveys per breeding season along a fixed survey route of typially 4-6 km in each square (Schmid et al., 2004). The Atlas survey data used here included 2318 sites, each of which was sampled in one year within each five-year period, while the MHB survey included 267 sites sampled annually. Further, we used land cover and bioclimatic data as environmental predictors. Land cover was represented with the CORINE Land Cover (European Union, 2022) classification (44 classes), obtained for the years 2000, 2006, 2012, and 2018 at a spatial resolution of 100 m. Climate was represented by the nineteen WorldClim bioclimatic variables. We used averaged annual values from the time period 1979–2013 with a spatial resolution of 30 arcsec (≈ 1 km) obtained from CHELSA Bioclim v1.2 (Karger et al., 2017; Karger et al., 2018). An overview of all data sources and their use in the modelling process is given in Appendix S1: Tables S1 to S3.

### 2.2 Modelling

Our dSDM comprised two components which are detailed in the following section: (1) a static habitat model that describes the habitat suitability in each year over the study region, and (2) a mechanistic individual-based model (IBM; Railsback & Grimm, 2019) that describes the population and range dynamics. IBMs use a bottom-up approach in which key processes are formulated at the individual level and are scaled up to the population level by numerical simulation (DeAngelis & Mooij, 2005). All analyses were conducted using the statistical programming language R (R Core Team, 2020) and several R packages (see below).

#### 2.2.1 Habitat suitability of the Swiss landscape

Habitat suitability was derived from a cSDM based on presence-absence data from the second atlas (2013-2016) data set. Because the red kite has been expanding its range in Switzerland during the past 50 years, this most recent data best reflects the underlying habitat requirements. For the cSDM, we assumed that most suitable habitats are already occupied even though they may not have reached their potential capacities yet. The red kite is a generalist and opportunistic raptor that breeds in a wide range of climates and habitats. Typical nesting habitat consists of forest patches with suitable roosting sites and adjacent open areas like grassland or agricultural fields that provide prey, which mostly consists of small mammals. Food sources like open waste dumps and carrion are readily exploited where present. To represent the availability of resources relevant for the occurrence of red kites, we used the CORINE 2012 land cover data and aggregated its 44 classes to seven land cover types (Appendix S1: Table S2). To represent climatic influences, all nineteen Bioclim variables were included (Appendix S1: Table S3). Since the red kite requires different habitats for nesting and foraging, it is a mobile species and occupies breeding home range sizes of about 4 to 5 km^2^ for males (Baucks, 2018; Nachtigall, 2008). To allow the cSDM to consider the diversity of habitat types, we used a grid cell size of 4 km^2^. Since the IBM was based on the same grid, this also constitutes a trade-off between the abilities to resolve both the effects of density-dependence on the one hand and dispersal displacements on the other hand. The high-resolution land cover data was aggregated to the target resolution of 2 km by calculating the proportional land cover in each cell. The bioclimatic data was coarsened to the 2 km-resolution by bi-linear interpolation between the grid cells.

To fit the cSDM habitat model, we first selected predictors from all land-cover and bioclimatic variables based on their univariate importance (assessed by the AIC of linear models with second-order polynomials) under the constraint that pairwise Spearman correlations must not exceed 0.7 (Dormann et al., 2013). The variables selected are labelled with an asterisk in Appendix S1: Tables S2 and S3. We then created an ensemble cSDM by taking the mean occurrence probability predicted by four different algorithms: binomial linear model with second-order polynomials and step-wise variable selection; binomial additive model with splines; random forest; and boosted regression trees. The predicted probabilities are subsequently interpreted as a habitat suitability index (HSI). The ensemble cSDM was then projected to Switzerland and a 12 km buffer around its border in the years 2000, 2006, 2012, and 2018, with varying land-cover data and constant bioclimatic variables. Climate was kept constant because it was considered only a minor driver of change in resource availability over the study period. The buffer was applied to reduce potential boundary effects in the IBM simulations. It was large enough to capture most dispersal events in the Swiss population. For all other years in the period of 1999-2019, the HSI values were linearly interpolated. To distinguish between habitat and non-habitat cells, we derived a binarisation threshold (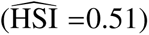) as the value yielding equal sensitivity and specificity (≈ 90%) and considered all cells with lower HSI values non-habitat.

#### 2.2.2 Individual-based model

We used the R package RangeShiftR (Malchow et al., 2021), which is an interface to the individual-based modelling platform RangeShifter 2.0 (Bocedi et al., 2021), to construct a dSDM based on the gridded habitat suitability maps described above. Next, we describe the main steps of the direct parameterisation, and refer to the full Overview, Design concepts and Details (ODD) protocol (Grimm et al., 2020) in Appendix S2 for all details. RangeShiftR explicitly simulates demography and dispersal in discrete unit-time steps, which here comprise one year. During each year, the processes “reproduction”, “juvenile dispersal”, “survival”, “development”, and “aging” are evaluated in this order for all individuals. The prior distributions on the respective process parameters (Table 1) were informed by literature data (Table S1) and expert knowledge. They were then updated with information contained in the survey data via Bayesian inference as described below.

**Table 1:** Process parameters of the IBM that were included in the Bayesian calibration and the parameters of their truncated normal prior distributions.

| Parameter name | Lower bound | Mean | SD | Upper bound |
| --- | --- | --- | --- | --- |
| Density-dependence $b^{-1}$ | 0.001 | 0.006 | 0.0025 | 0.020 |
| Fecundity $\phi_0$ | 0.5 | 1.66 | 0.51 | 5.0 |
| Survival prob. $\sigma_1$ | 0.01 | 0.42 | 0.08 | 0.99 |
| Survival prob. $\sigma_2$ | 0.01 | 0.68 | 0.09 | 0.99 |
| Survival prob. $\sigma_3$ | 0.01 | 0.80 | 0.05 | 0.99 |
| Development prob. $\gamma_1$ | 0.01 | 0.80 | 0.10 | 0.99 |
| Development prob. $\gamma_2$ | 0.01 | 0.55 | 0.10 | 0.99 |
| Emigration prob. $e_1$ | 0.01 | 0.80 | 0.10 | 0.99 |
| Settlement inflection pt. $\beta_s$ | -15.0 | 4.0 | 4.0 | 15.0 |
| Dispersion parameter $\nu$ | 1.0 | 50.0 | 250.0 | 500.0 |

Our model is female-based, since females primarily determine the population dynamics in red kites. Their development is described in three stages (Fig. 2), with classifications and age ranges adopted from Sergio et al. (2021) and Newton et al. (1989): Dispersing juveniles are one or two years old, sub-adults establish a territory within their second to sixth year, and breeding adults are aged between three and twelve years. A senescent stage was not included in the model because it does not contribute to the overall fecundity and non-breeding adults are not monitored in the survey. These age limits are not strict, as the stage transitions are modelled probabilistically (Fig. 2). The transition probabilities *τ*_*m*,*n*_ are expressed as survival probabilities of stage *s*, *σ*_*s*_ = *τ*_*s*,*s*_ + *τ*_*s*,*s*+1_, and the development probabilities *γ*_*s*_ = *τ*_*s*,*s*+1_ *σ*^−1^. Both can independently vary between zero and one. The development probabilities are assumed as *γ*_1_ = 0.80 ± 0.10 for stage 1 and *γ*_2_ = 0.55 ± 0.10 for stage 2, to approximately yield the described age classes (Appendix S1: Fig. S1). The survival probabilities *σ* are taken from Katzenberger et al. (2019) for all three stages: *σ*_1_ = 0.42 ± 0.08 and *σ*_2_ = 0.68 ± 0.09 and *σ*_3_ = 0.80 ± 0.05, which is also in accordance with Schaub (2012) and Newton et al. (1989).

**Figure 2:**
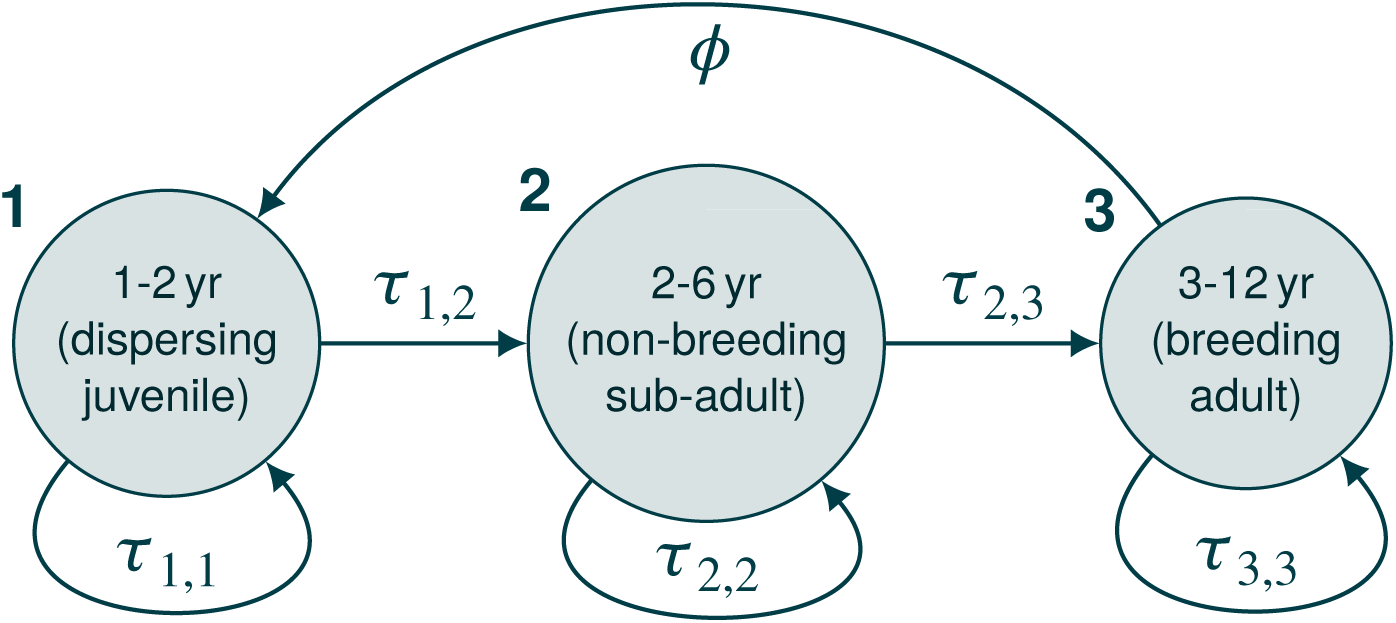
Life-cycle graph of the population model for the red kite with three developmental stages. The probability for an individual in stage *s* to stay in its stage over one time step (1 year) is denoted by *τ*_*s*,*s*_, and to move to the next stage is denoted *τ*_*s*,*s*+1_. Only stage 3 produces offspring, with a fecundity of *φ*.

Fecundity *φ* was assumed to be density-dependent and was modelled as exponentially decaying with population density. Each cell *i* is characterised by a local strength of demographic density-dependence *b*_*i*_, which is obtained as the global strength *b* divided by the cell habitat suitability HSI_*i*_, *b*_*i*_ = *b* HSI^−1^, given in units of cell area (*a*_*c*_ = 4 km^2^). Fecundity follows the relation *φ*_*i*_(*n*_*i*_) = *φ*_0_ e^−*bi ni*^, where *n*_*i*_ denotes the density of adults in stages 2 and 3 in cell *i* (i.e. juveniles do not contribute towards this density-dependence). The base value *φ*_0_ is the required process parameter and denotes the theoretical fecundity at zero population density. Nägeli et al. (2021) report a realised fecundity of 1.77 ± 0.70, which agrees with Schaub (2012) and Nachtigall (2008). We assumed that this value is reached at a density of 25 breeding pairs (BP) per 100 km^2^ (i.e. 1 BP per cell) and halved it for our female-only model: 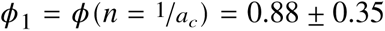. We can then get *φ*_0_ from *φ*_0_(*b*) = *φ*_1_ e^*b*/HSI^, with *b* as a calibration parameter that controls the strength of density-dependence in fecundity. The HSI over all habitat cells had a mean and standard deviation of 80% ± 12% and we assumed that the lower and higher fecundities were attained in the lower and higher quality habitats. This was given with a range of *b* = (0.50 ± 0.15) *a*_*c*_ (Appendix S1: Fig. S2), yielding *φ*_0_ = 1.65 ± 0.51.

Dispersal is explicitly modelled in three stages: emigration, transfer and settlement (Travis et al., 2012). Red kites are strongly philopatric (Newton et al., 1989) so that emigration was modelled as occurring in the first developmental stage, i.e. dispersing juvenile, only. The emigration probability was assumed constant at *e*_1_ = 0.64 ± 0.10, meaning that an expected proportion of 87% of juveniles have dispersed within their first two years. This value best matched observations in which 42% of females of a cohort have emigrated after year one and 45% after year two (own unpublished data), suggesting a larger proportion of emigrants among two-year-old juveniles than among one-year-old juveniles. The transfer phase described the movement of a dispersing individual through the landscape. It was modelled as a strongly correlated random walk in a random direction with a step length equal to the cell size. After each step, the option to end the movement and settle in a cell for future breeding site was evaluated. Settlement was only possible in habitat cells and its probability was density-dependent with a sigmoid relationship (Appendix S1: Fig. S3). Its inflection point *β*_*s*_ was a calibrated parameter and was estimated as 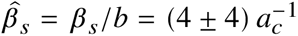. The maximum settlement probability and the slope parameter were fixed at *⍺*_*s*_ = −1 and *s*_0_ = 0.75, respectively, in order to reduce flexibility. At these values, the choice of just *β*_*s*_ allows to tune the density-dependent settlement across a wide range of reasonable relationships (Appendix S1: Fig. S3). The maximum number of steps in the random walk was set to 10. Therefore, depending on the availability of sparsely populated habitat, individuals exhibit dispersal distances between 2 and maximal 20 kilometres, which is consistent with observations (own unpublished data; Newton et al., 1989; Nachtigall, 2008). Longer-range dispersal events are also frequently observed, but excluded from the model due to the small study region and the mountainous terrain. Immigration or emigration of individuals across the system boundaries was not considered.

The initial conditions of each simulation were stochastic. The number of adult individuals in each cell was drawn from a Poisson distribution whose mean values were predicted from a generalised linear model. This model of the initial red kite distribution was an autoregressive distribution model (Dormann et al., 2007) of the earlier atlas data (1993-1996) with the spatially interpolated values of atlas counts as its sole predictor (Appendix S1: Fig. S4). The number of juveniles and sub-adults was subsequently estimated from the demographic rates under the assumption of a stable stage distribution.

### 2.3 Bayesian calibration

We used Bayesian inference to evaluate the joint posterior distribution of nine model parameters *θ* based on their prior distributions *p*(*θ*) and the likelihood *l*(*θ*) (Fig. 1). The priors express the a-priori information that we assumed about likely parameter values as summarised in Table 1. The likelihood function *l* (*θ*) measures the fit of a model *M*, parameterised with *θ*, to the monitoring data. The estimated model parameters are: the strength of density-dependence 1/*b*; six demographic probabilities for survival of all stages (*σ*_1_, *σ*_2_, *σ*_3_), juvenile and sub-adult development (*γ*_1_, *γ*_2_), and the base adult fecundity (*φ*_0_); as well as two dispersal parameters to control the emigration (*e*_1_) and settlement probabilities (*β*_*s*_). Additionally, we calibrated a dispersion parameter *v*, which is introduced below.

As priors *p*(*θ*) we chose as truncated normal distributions for all parameters. Their means and standard deviations were informed from the literature and expert opinion and they were bounded to their respective valid parameter ranges (Table 1). As calibration data, we used observed abundances from the MHB survey, *D*_MHB_. Based on this data, we defined a likelihood *l* (*θ*) = *p*(*D*_MHB_ | *θ*, *M*) as follows: For a given parameter vector *θ*, the RangeShiftR simulation model (*M*) is run and the output abundance data are aggregated and averaged over twenty replicate runs of the model. The result *D*_sim_ was compared with the MHB counts under the assumption of a negative-binomial error distribution (NB), so that for an observation at site i and time t,

*l* _*i*,*t*_ (*θ*) = *Pr*_NB_(*D*_MHB,i,t_ | *μ*=*D*_sim,i,t_, *v*). The parameter *ν* describes the error over-dispersion and was also estimated from the data. It arises in an alternative formulation of the negative-binomial probability mass function *Pr*_NB_ formulated in terms of its mean *μ* and dispersion *ν*, instead of the more common success probability 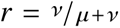 and the target number of successes *n* = *v*. Therefore, its variance is given by 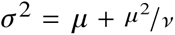. It approaches *μ* from above when *v*→ ∞, as the negative binomial converges to the Poisson distribution. The amount of over-dispersion can thus be tuned by the value of *v*, rendering *Pr*_NB_ an appropriate error description for potentially over-dispersed count data.

Due to the stochasticity inherent in our simulation model, the likelihood values calculated from repeat simulations were still stochastic to some extent. They thus represent an estimator of the exact likelihood. Conceptually, this not a problem for our Markov chain Monte Carlo approach (MCMC, details below), since the pseudo-marginal theorem guarantees that the MCMC sample still converges to the exact posterior distribution (Andrieu & G. O. Roberts, 2009). In practice, however, large variances in the likelihood estimator can increase the time required for MCMC convergence dramatically when the sampler gets stuck at occasional high values. To reduce the variance in the likelihood estimates, we aggregated the abundance data within spatio-temporal blocks of 14 ×14 grid cells in space and three years in time (Appendix S1: Figs. S5 and S6). These aggregation factors were chosen with the target of reaching a variance below ten on the logarithmic scale in repeated likelihood evaluations for a given *θ* (Appendix S1: Fig. S7). The aggregation resulted in 57 spatial and 5 temporal blocks, within which the observed and simulated red kite densities were compared. Under the usual independence assumptions, the total likelihood was then expressed as the product over all such blocks: 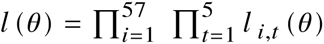.

To validate the calibration setup and assess the sensitivity of the likelihood *l* (*θ*) to changes in the model parameters *θ*, we performed a local sensitivity analysis (Appendix S1: Fig. S7). For this, a test data set *D*_SA_ was simulated from the model with all parameters at their mean prior values. Then, one parameter at a time was varied within the boundaries of its prior distribution while keeping all other parameters at their mean and estimating the likelihood with respect to *D*_SA_. Further, we performed a global sensitivity analysis with Morris’ elementary effects screening method (Morris, 1991).

To estimate the joint posterior distribution based on the defined *p*(*θ*) and *l* (*θ*), we used a MCMC sampling scheme (Luengo et al., 2020). Therein, the posterior density *p*(*θ* | *D*_MHB_, *M*) of a series of given parameter sets *θ* is evaluated according to Bayes’ rule. The utilised MCMC algorithm was a variant of the adaptive Metropolis sampler, namely the differential evolution sampler with snooker update (DEzs, Braak & Vrugt, 2008), as implemented in the BayesianTools R package (Hartig et al., 2019). Every calibration run included three independent DEzs-MCMC instances with a length of 2 × 10^5^ iterations, of which the first 5 × 10^4^ were discarded as a burn-in. Each DEzs, in turn, consisted of three inter-dependent internal chains, so that each calibration comprised a total of nine chains. The chains were checked for convergence using trace plots, trace rank plots (Vehtari et al., 2021) and the multivariate potential scale reduction factor (psrf; Gelman & Rubin (1992)). A chain was considered approximately converged if its multivariate psrf value had dropped below 1.10 and the trace and trace rank plots showed well mixed chains.

To assess the information gained in the calibration, the sampled posterior distributions were contrasted with the prior distribution. To this end, the parameter estimates were compared with respect to the medians and quantiles of their respective marginal distributions. To evaluate if and by how much the uncertainty was reduced, we assessed and compared the distribution breadth by calculating the width of the highest-posterior-density intervals (HPDIs).

### 2.4 Cross-validation and prediction

We employed a spatial block cross-validation scheme to evaluate the model fit without duplicate use of data for both model calibration and validation (D. R. Roberts et al., 2017). To this end, the data was split into five spatially contiguous folds (Fig. 1 and Appendix S1: Fig. S5). For each fold, the respective subset of MHB data was held out and the model was fit to the remainder of the data. To ensure that the folds covered largely identical spaces of environmental conditions, we chose longitudinally structured folds that include a similar altitudinal profile. For model validation, the respective calibration results for each fold were used. For final model projections, in turn, a separate calibration on the full data set was used.

Posterior model predictions were generated by taking a sample of 1000 draws from the joint posterior, running the dSDM with each drawn parameter vector, and calculating the mean, median and 95%-credibility interval (CI) of the simulated abundances. Prior predictions were obtained in the same way but using draws from the prior distribution. Both prior and posterior predictions were run for the time covered by the MHB data and additional 30 years forward with constant habitat suitabilities, i.e. no changes in land cover or climate were considered. This projection provides an estimate of the potential current population size and distribution, without making a prediction to future conditions.

To assess the model’s predictive performance, we calculated Harrell’s c-index (Newson, 2006, using the function *rcorr.cens* from the Hmisc R package), a rank correlation index that generalises the AUC index to non-binary response variables. It quantifies the probability that for a given pair of data points the ranking of predictions matches the ranking of observations. This measure was used in Briscoe et al. (2021) as a form of temporal AUC to assess the fit to temporal trends. We use it here as an index that is applicable to abundance projections and can be interpreted like the AUC for occurrence projections.

## 3 Results

### 3.1 Sensitivity analysis

Based on the local and global sensitivity analyses (Appendix S1: Fig. S7 and Fig. S8), we found that the likelihood estimates responded most strongly to variation in the strength of density-dependence 1/*b*, the adult base fecundity *φ*_0_ and the three survival probabilities *σ*_1_, *σ*_2_, and *σ*_3_. Therefore, we expected that these parameters will calibrate best under our setup, while the development probabilities *γ*_1_, *γ*_2_, and the emigration probability *e*_1_ would be only weakly informed by our survey data through the specified likelihood.

### 3.2 Model calibration

To calibrate the model parameters, we ran independent DEzs-MCMC chains on different training data sets: five chains were run on the separate folds of the cross validation and one on the full data set. Differences between the respective sampled posterior distributions can therefore arise both because of differing convergence and because of the data selection. We find that all posteriors converged roughly in the same area of the parameter space, as the variance over the five folds was small. Their marginal distributions take on similar medians and quantiles.

A comparison between the prior and posterior distributions revealed how the consideration of the MHB survey data informs the initial parameter estimates that were obtained directly from literature data. Notably, the medians of the marginal distributions for fecundity *φ*_0_ and the survival probabilities of the first and second stage, *σ*_1_ and *σ*_2_, have shifted significantly, whereas those of the other parameters remained largely unchanged. The marginal posterior distributions for each parameter and each spatial fold are represented by box plots in Fig. 3, and those for the calibration to the full data set are shown in Appendix S1: Fig. S12. Comparing the HPDIs of the prior and posterior distributions, we found substantially narrower posteriors and thus reduced uncertainty for the strength of density dependence 1/*b*, fecundity *φ*_0_ and the survival probabilities of stages one and two, *σ*_1_ and *σ*_2_. These parameters had already responded strongly in the sensitivity analysis. No uncertainty reduction nor a significant change in point estimate were found for adult survival *σ*_3_ (Appendix S1: Fig. S13). This was contrary to our expectation based on the sensitivity analysis, but this parameter already had the most informative priors to begin with. The dispersion parameter of the negative binomial error model was calibrated to a very large value, yielding a variance that was close to that of a Poisson distribution. Thus, only slight over-dispersion was detected relative to a Poisson-distributed error.

**Figure 3:**
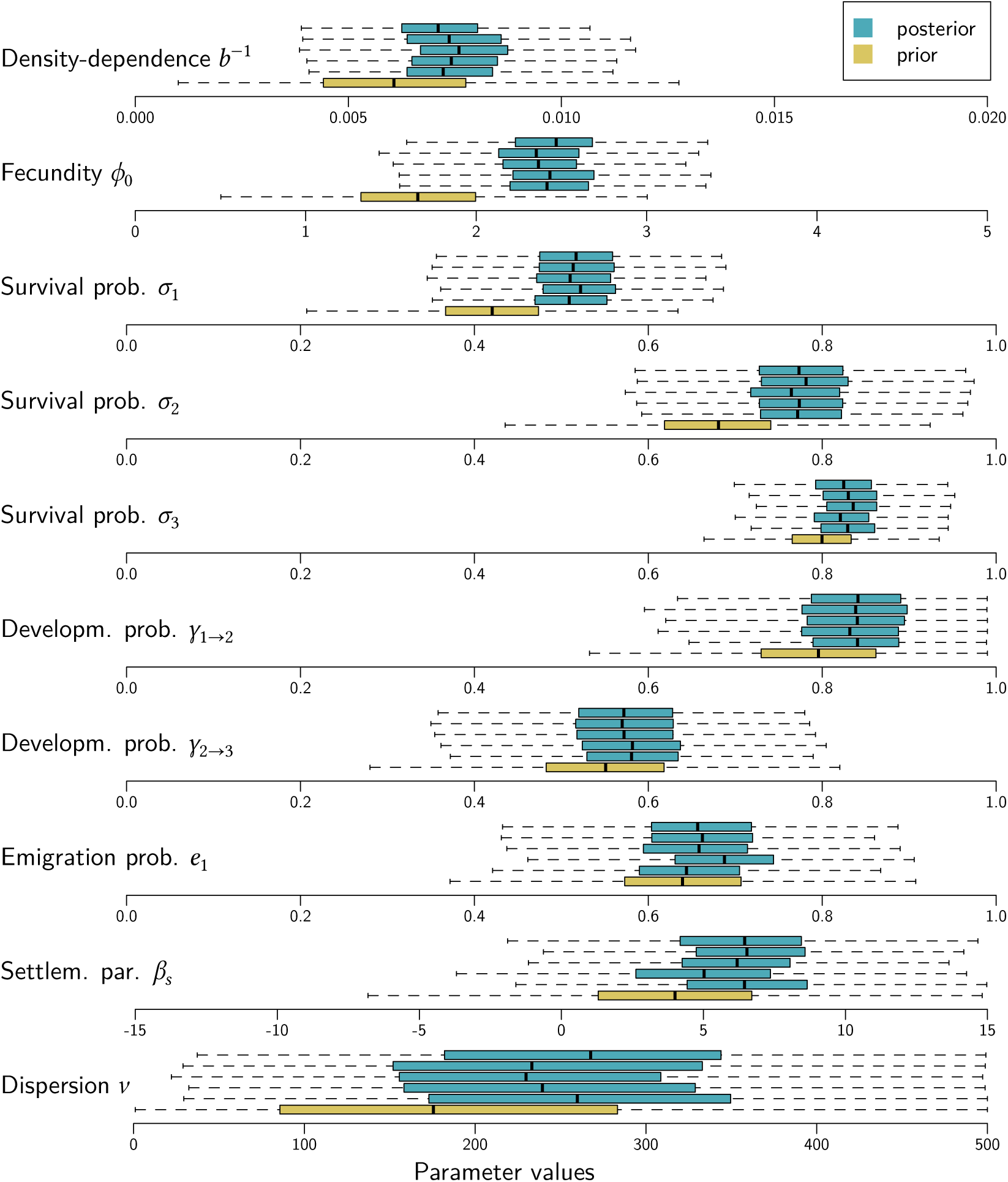
Box plots summarise the marginal prior (yellow) and posterior (blue) distribution for each calibration parameter and for all five spatial folds. The black bar marks the median, the boxes show the inter-quartile range, the whiskers extend to the most extreme data point which is no further away from the box than 1.5 times its length.

The convergence of all DEzs-MCMCs was regarded sufficient, based on the conducted diagnostics. However, there were considerable differences between the folds due to the varying number of MHB sites included: The chains reached multivariate psrf values of 1.05, 1.02, 1.05, 1.09, and 1.04, respectively, for folds 1 to 5. Convergence was further assessed using trace plots (Appendix S1: Fig. S9), trace rank plots (Appendix S1: Fig. S10) and psrf plots (Appendix S1: Fig. S11), which were all satisfactory. No substantial correlations between the parameters were detected (Appendix S1: Fig. S14).

### 3.3 Model validation

The spatial-block cross-validation was evaluated by calculating the c-index per spatial fold and for different subsets of the MHB data (Table 2). First, it was calculated over all observations, i.e. all site-year combinations within a fold, independently. The overall value of 0.88 indicates an excellent fit to the validation data. However, the results were quite variable across folds (see also Appendix S1: Fig. S15), which is likely due to the differing number and information content of the included MHB sites. Second, focusing on regional abundance dynamics, we calculated the c-index for the time series of the total abundance within each fold, consistently yielding excellent values between 0.92 and 0.94 (see also Appendix S1: Fig. S16). This confirms that averaging the abundance over large regions further increases the accuracy of temporal predictions. Third, we were interested in the performance of our dSDMs at those MHB sites that showed the highest variance in red kite counts, since highly fluctuating population sizes are often of special conservation interest but are usually harder to predict. To this end, we ranked all MHB sites by their count variance and computed the c-index over the top 15% most variable sites. The folds scored significantly lower, showing an overall value of 0.66 (see also Appendix S1: Fig. S17), which signifies a substantial drop in performance and indicates fair predictions for highly variable sites. Again, the different folds show very variable results that range from 0.59 to 0.75, depending on the specific sites they include.

**Table 2:** Evaluation of the spatially-blocked cross-validation (CV). Each fold was used as test data for a separate calibration, which used the remaining folds as training data. Predictions to the test data were evaluated using the c-index, given per row by its mean and standard deviation in brackets. The c-index was computed on different subsets of the MHB data, given per column: The spatio-temporal c-index compares each observation (site-year) independently, the temporal c-index compares time series of total abundance per fold, and the last column gives the c-index over the sites with the 15% highest variance in local population size.

| Spatial CV fold | Spatio-temporal | Temporal | High variance |
| --- | --- | --- | --- |
| Fold 1 | 0.69 (0.05) | 0.94 (0.05) | 0.67 (0.11) |
| Fold 2 | 0.86 (0.02) | 0.93 (0.05) | 0.59 (0.06) |
| Fold 3 | 0.89 (0.02) | 0.94 (0.04) | 0.65 (0.07) |
| Fold 4 | 0.92 (0.01) | 0.92 (0.05) | 0.75 (0.05) |
| Fold 5 | 0.85 (0.04) | 0.92 (0.06) | 0.73 (0.07) |
| All folds | 0.88 (0.01) | 0.94 (0.04) | 0.66 (0.03) |

### 3.4 Model projections

The model was used to generate projections to places not covered by the MHB survey by simulating red kite abundance over the whole extent of Switzerland. This allows to compare these projections to the Swiss breeding bird index, which estimates the total population trend relative to the year 1999 (Knaus et al., 2022) and thus offers an additional source of validation data. Further, by running the model forward beyond the MHB data period and under stable environmental conditions, we estimated the size and range of the current potential population.

Prior and posterior predictions of total red kite abundance during the entire survey period and thirty years onward, assuming constant habitat suitabilities, are shown in Fig. 4. The posterior predictions show very good fit to the Swiss breeding bird index. Comparing the prior and posterior predictions of our model gives more evidence that the calibration was able to gain substantial information from the survey data: the model fit was improved considerably and output uncertainty was reduced.

**Figure 4:**
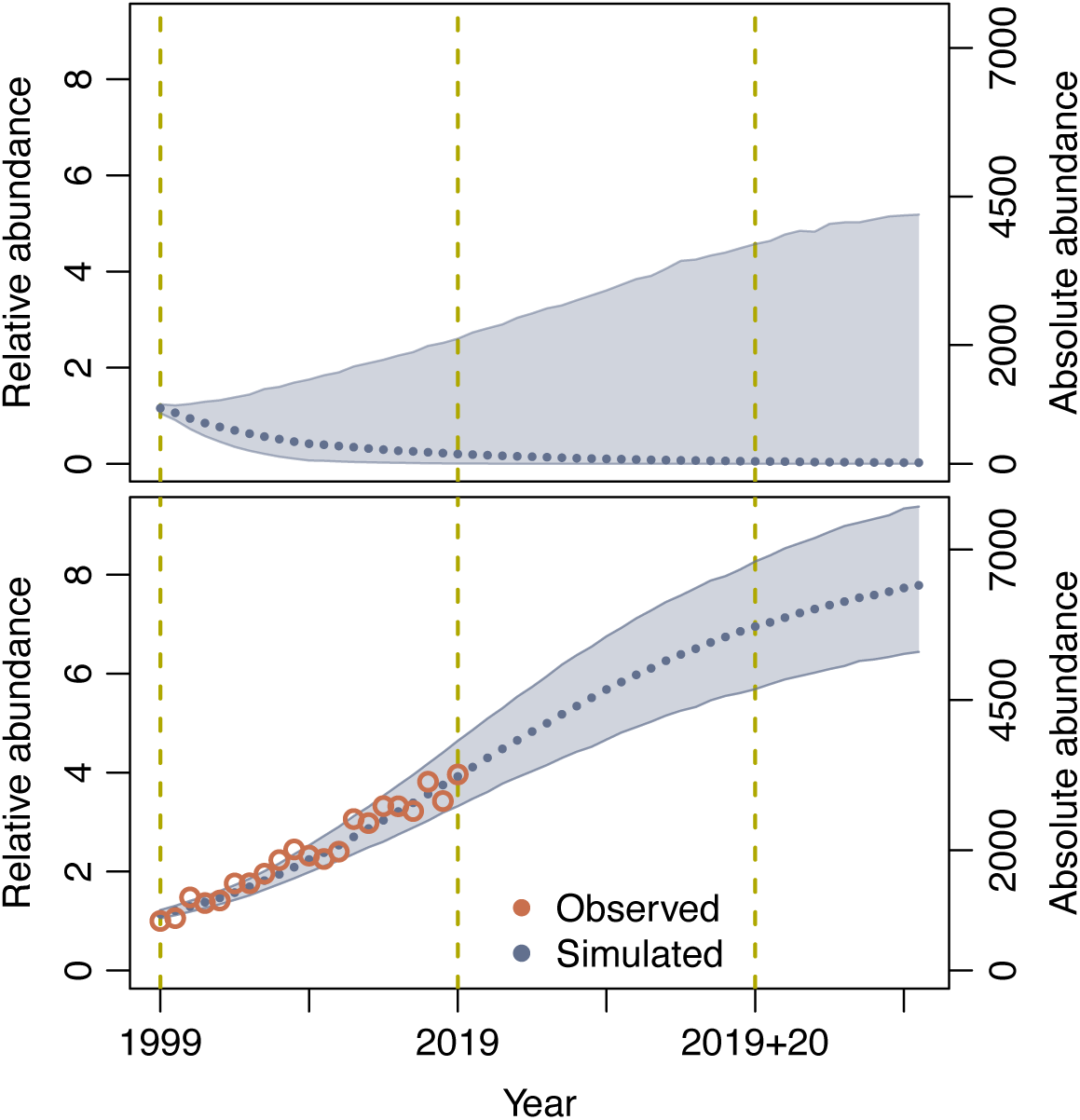
Prior (top) and posterior (bottom) simulations of the abundance time series of red kite in Switzerland. The blue line and band show the median and 95%-credibility interval of total number of predicted breeding pairs (#BP), with relative values (with respect to year 1999) on the left and absolute values on the right y-axis. The bottom panel shows the cross-validated posterior predictions together with the breeding bird index (red circles, relative y-axis only) for comparison. The dashed vertical lines mark the years for which spatial projections are depicted in Fig. 5. After the last year of survey data, 2019, the environmental conditions are kept constant.

Simulations from the prior show negative population trends in most cases, and they have a large 95%-CI that includes predictions of 5 to 2100 breeding pairs in year 2019. The posterior predictions, in contrast, show increasing trends throughout, and a much narrower CI. They exhibit a comparably steep increase in abundance over the past twenty years, in accordance with the strong increases in red kite abundance that were recorded during this time. Forward simulation shows that today’s potential equilibrium population size amounts to 6400 (95%-CI: 5300-7700) breeding pairs.

Spatial projections were made to the whole country as three snapshots in time (Fig. 5): at the beginning (1999) and end (2019) of the survey data set, as well as after a continuation of further twenty years (2019+20). These projections mirror the rapid range expansion of red kite range that Switzerland has seen in the past two decades. However, the continuation shows a relatively stable range with increasing population densities, suggesting that the current population has not yet reached the carrying capacity in all colonised areas. The same maps were created from prior predictions for comparison (Appendix S1: Fig. S19). They exhibit a contracting range over time, that deviates substantially from the posterior maps, again indicating the effective inclusion of information from the MHB data.

**Figure 5:**
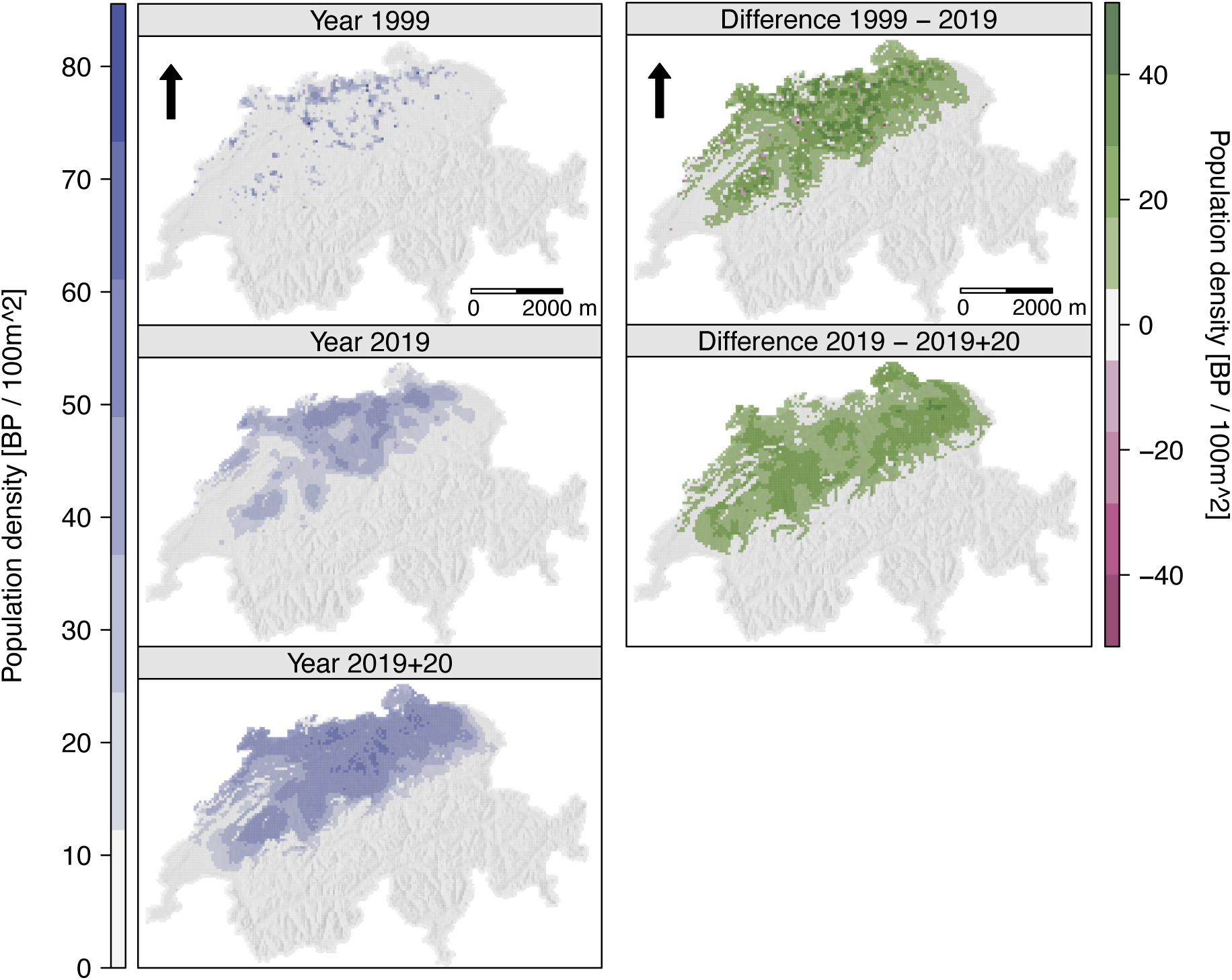
Mean posterior predicted population densities for the years 1999 and 2019, and after 20 further years under constant conditions (left column) as well as the differences between those years (right column). The arrows indicate north.

This comparison of prior and posterior distributions and their respective predictions can shed light on the main drivers of the presented results. While the prior predictions exhibit a tendency towards decreasing populations and contracting ranges, the posterior predictions reproduce the observed patterns closely. Comparing the marginal distributions (Fig. 3), the main drivers of these disparate predictions appear to be fecundity and early survival rates. They responded most strongly to the information incorporated from the MHB data via the Bayesian calibration. Taken together, these three influential parameters suggest that reproductive success was determined to play a key role in driving the resulting increases in local density and distribution. In contrast, changes in habitat suitability over the study period seem to have had a lesser effect on the resulting population. This was assessed in a simple analysis of the sensitivity of simulated abundance to habitat suitability. We compared the abundance time series from Fig. 4 with two counter-factual scenarios in which the habitat suitabilities of each year were raised or reduced by five (out of 100) habitat suitability points (Appendix S1: Fig. S18). By the last year of MHB data, 2019, this intervention had an effect of 9-10% on total abundance, which is small compared to the effect of the calibration (Fig. 4). We thus conclude that the population increases are not predominantly driven by a changing environment, but by transient dynamics to an equilibrium with much higher population size.

## 4 Discussion

Reliable methodology for understanding and predicting a species’ population and range dynamics will be crucial to inform decision making in the future. Dynamic, spatially-explicit, process-based distribution models (dSDMs) provide valuable advances towards improved biodiversity forecasts (Urban et al., 2016) but are currently underused due to technical and data challenges and limited guidance for applications (Briscoe et al., 2019; Zurell et al., 2022). This study contributes to overcoming these challenges. We demonstrated the practicability of a complete modelling workflow for dSDMs with a case study of a conservation-relevant population, the red kite in Switzerland. This included calibrating a complex stochastic simulation model to heterogeneous empirical data, interpreting the results, and validating the model by cross-validation. Thanks to the use of Bayesian inference, we can integrate direct and indirect knowledge on the process parameters, account for their uncertainty and propagate it to model projections. Our model captures the Swiss red kite population trends with higher spatial and temporal predictive accuracy than achieved with correlative models in a previous study (details below, Briscoe et al., 2021). The model suggests that the potential population size under current environmental conditions is much larger than presently realised and that this may be a result of the population’s history. The workflow exemplified here can be readily adapted to other species, if an adequate model, prior parameter estimates, as well as response data (e.g. occupancy or abundance data) are available, and it promises to yield improved parameter estimates and more accurate, validated model projections.

The process-based dSDM used here to demonstrate our workflow was built with the individual-based modeling platform RangeShiftR (Malchow et al., 2021). It explicitly considers relevant ecological processes such as demography and dispersal and includes crucial mechanisms such as density-dependence. Therefore, transient dynamics, that arise when a distribution is not in equilibrium with its environment, can be reproduced and dynamic responses to change can be represented. The IBM structure was determined based on expert opinion and the direct (prior) parametrisations of the process parameters were derived from literature data. IBM approaches are particularly suited for the direct estimation of their model parameters (e.g. survival probability or dispersal distance) because they formulate all processes from the perspective of the individual (Railsback & Grimm, 2019), where they can be estimated from data obtained in observational studies (e.g. mark-recapture). The prior estimates were updated using the MHB abundance data within a Bayesian inference. Here, IBMs have the advantage of realistically modelling small local populations of a few individuals by incorporating demographic stochasticity, so that the survey data can be used at a high spatial resolution. Depending on the research question and the available data, however, a different model formulation may be more adequate, such as spatially-explicit population-based models (implemented for example in the R package steps, Visintin et al., 2020). For a successful parameterisation using the presented framework, certain data requirements should be met: the utilised model should have model parameters whose priors can be informed by ecological theory and direct estimates from field and experimental measurements, and it should generate outputs that can be compared to observational data via a plausible error model that can be expressed as a likelihood function.

The successful calibration of parameters in process-based dSDMs can produce new insights, since they have a well-defined ecological meaning. Comparing the prior and posterior distributions of our model, we found that some parameters in particular, e.g. the adult fecundity *φ*_0_, its density-dependence 1/*b*, and the survival probabilities of juveniles and sub-adults, *σ*_1_ and *σ*_2_, responded strongly to the inclusion of additional information from survey data. This behaviour was predicted well from the local and global sensitivity analyses. The detected sensitivity further indicates that the model projections are responsive to these parameters, thereby suggesting potential pathways for conservation measures, e.g. highlighting the protection of young individuals and supporting nest success. Pfeiffer & Schaub (2023) reached the same conclusion and identified the breeding output and adult survival as the main demographic drivers when modelling the German red kite population with an integrated population model. Further, their estimates of stage-wise survival probabilities are in strong agreement with ours. The prior estimate of 1/*b* was corroborated and those of *φ*_0_, *σ*_1_ and *σ*_2_ were corrected to higher values, while the uncertainty around all four estimates was reduced. These corrections in parameter distributions also drive the better fit of the calibrated model projections to the data as well as the reduced output uncertainty. Interestingly, it was shown that the calibration could gain information even on the early developmental stages (1 and 2) that were not recorded in the calibration data, which held abundances of stage 3 only. This is facilitated by the ecological assumptions that are imposed by the model structure which discerns the stages by their ability to disperse (only stage 1) or reproduce (only stage 3).

The discrepancies found between prior and posterior parameter estimates are driven by different factors (Cailleret et al., 2020): Firstly, there can be a true difference, for example, if the prior estimates were obtained from different study populations. In our case, the prior fecundity was based on a measurement from a Swiss sub-population with a high breeding density, which may have a lower fecundity than the Swiss average. The prior survival probabilities are taken from a German red kite population, that shows a slightly negative population trend, and the upward correction seems to better match the increasing Swiss population. Red kite in Switzerland benefit from public feeding (Cereghetti et al., 2019) and many sub-adult individuals change from migratory to resident behaviour, which can also increase survival. Secondly, an important source of deviations between empirical and calibrated parameter estimates is model error. Our IBM captured only a subset of the multiple eco-evolutionary processes that underlie the observed abundance patterns. Therefore, the calibrated parameters will account for missing processes to some extent. This highlights the need for further development of dSDMs to include more mechanisms and thus to fit observed data more closely. It further emphasises the importance of effective integration of direct and inverse calibration to estimate parameter values and their uncertainties, since projections and derived management decisions can be highly sensitive to the final parameterisation.

The validation of model predictions to assess model performance is common practice in the application of cSDMs (Sillero et al., 2021), but is often missing with dSDMs. With the presented workflow, we have successfully applied spatial-block cross-validation to a dSDMs by calibrating the model to each of five spatially contiguous regions within the study area. By spatially blocking the hold-out data we reduced the amount of spatial autocorrelation between the training and testing data, which is often present in abundance data and only insufficiently reduced by other cross-validation schemes such as randomised leave-p-out. This yields a more realistic assessment of predictive performance for interpolation. For other types of data, different blocking techniques may be more adequate (D. R. Roberts et al., 2017). The folds were selected carefully in a way that the same range of environmental conditions is represented in each one, so that model evaluation does not involve extrapolation to new environmental conditions. Performing a cross-validation is usually computationally expensive, as the calibration needs to be repeated for each set of hold-out data. Therefore, a suited validation method has to be chosen carefully. Alternatives include approximation to leave-one-out validation by WAIC (Vehtari et al., 2017).

The full red kite dSDM was evaluated based on its cross-validated abundance predictions using Harrell’s c-index as a measure of predictive performance, which indicated excellent predictive accuracy (c-index: 0.88). In sites with highly fluctuating abundances performance dropped considerably, and only provided fair predictions (c-index: 0.66). A similar analysis was conducted by Briscoe et al. (2021) who compared the accuracy of correlative SDMs and dynamic occupancy models (Kéry et al., 2013) that were fitted to the MHB data of 69 Swiss birds, including the red kite.

They found that predictive ability of occupancy was high for all examined model types when assessed across all sites (mean AUC > 0.8) but much lower when specifically testing only sites that showed occupancy change (mean AUC 0.64-0.71). The AUC metric is based on predictions of occupancy only and therefore generally scores above the c-index, which ranks abundances. Collapsing our abundance predictions to occupancy and comparing them to Briscoe et al. (2021) in terms of the AUC, our calibrated IBM surpasses the mean of their red kite SDMs, both across all sites (mean AUC of 0.91 versus 0.85) and occupancy-switching sites only (0.80 versus 0.67). Especially in range shifting populations, such as the red kite in Switzerland, process-explicit dSDMs can outperform correlative approaches because they make no equilibrium assumption. Our model also showed clear advantages over the dynamic occupancy models in Briscoe et al. (2021), likely due to the explicit consideration of spatially-explicit processes such as density-dependence in population dynamics and dispersal.

The calibrated model was run forward under current environmental conditions in order to explore the potential population size and distribution. In the same way, it could be used to assess population trends under certain scenarios such as conservation measures involving habitat improvements or regulation of demographic rates. For predictions of future population dynamics, however, expected changes in land use and climate have to be taken into account. In our dSDM, these variables are only considered through the cSDM-derived, scalar habitat suitability and thus can not impact demographic processes directly and independently, as ecological theory suggests. More complexity and mechanistic understanding could be incorporated by substituting the habitat model with direct relationships of species traits like demographic rates with environmental variables. Such a demographic range model is adequate for predictions under climate change (Schurr et al., 2012; Malchow et al., 2023). As a further limitation, the habitat map that underlies our model consists of a cSDM fitted to recent Atlas data. It is possible that suitable but not yet occupied parts of the red kite niche were missed in this data and therefore the future range would be underestimated by the projections. This limitation can be circumvented by using a habitat model that does not rely on the equilibrium assumption, e.g. a rule-based model derived from knowledge about the species’ habitat requirements or an eco-physiological SDM (Kearney & Porter, 2009). Moreover, the observed increases in range and density may in part be fueled by individuals that were not recruited in the study region but immigrated from surrounding high-density populations that were not considered in the model. More potential for model improvement lies in implementing additional processes such as mating systems, species interactions, or genetic and behavioural adaptation. Their successful parameterisation, however, will require adequate data.

The inverse calibration of dSDMs from observational data is also possible with other methods like pattern-oriented modelling (POM; Grimm et al., 2005; Mortensen et al., 2021) or approximate Bayesian computation (ABC; Beaumont, 2010; Hauenstein et al., 2019), the latter of which has already been demonstrated in the RangeShifter model (Dominguez Almela et al., 2020). Independent of the chosen calibration method will the accurate parameterisation of dynamic and mechanistic SDMs remain a challenge until the paucity of high-quality ecological and monitoring data is alleviated (Oliver et al., 2012; Kissling et al., 2018). Which parameters of a dSDM can be successfully calibrated depends on the available calibration data. Here, the type of data collected within monitoring programs plays an important role as all model output quantities can principally be used for inverse parameterisation. In our case study, for example, the abundances of juveniles and sub-adults were output from the IBM but could not be used for calibration because age classes are not distinguished in MHB surveys. Generally, the high uncertainties in parameter estimates caused by data limitations translate to large credibility intervals in model projections, reducing the utility for conservation applications. Here, it is a clear advantage of the Bayesian framework that sources of outcome uncertainty are explicitly quantified and can thus be addressed, for example in targeted monitoring programs.

In conclusion, this case study shows how an individual-based dSDM can be built with RangeShiftR, calibrated using Bayesian inference, and validated by cross-validation. We demonstrated how the inclusion of monitoring data refined parameter estimates and greatly improved model fit and prediction accuracy, thereby offering improved insights into underlying mechanisms. Well calibrated and validated process-based models offer compelling advantages over the currently most common static models. They are able to inform science-based management decisions and the design of proactive conservation measures (Zurell et al., 2022). Future progress in this field should be directed towards developing more flexible and accessible modelling tools, assessing their data requirements for effective parameterisation, and validating them against independent targets.

## Supporting information

ODD protocol for IBMs

## Acknowledgements

AM and DZ were supported by Deutsche Forschungsgemeinschaft (DFG) under grant agreement No. ZU 361/1-1.

## Author Contributions

A.-K. Malchow: Conceptualization (equal), Formal Analysis (lead), Methodology (equal), Software (lead), Data Curation (equal), Visualization (lead), Writing – Original Draft Preparation (lead), Writing – Review & Editing (equal).

G. Fandos: Conceptualization (equal, Methodology (equal), Writing – Review & Editing (equal).

U. G. Kormann: Conceptualization (equal), Data Curation (equal), Methodology (equal), Writing – Review & Editing (equal).

M. Grüebler: Conceptualization (equal), Data Curation (equal), Methodology (equal), Writing – Review & Editing (equal).

M. Kéry: Conceptualization (equal), Data Curation (equal), Writing – Review & Editing (equal).

F. Hartig: Conceptualization (equal), Formal Analysis (support), Methodology (equal), Supervision (support), Writing – Review & Editing (equal).

D. Zurell: Conceptualization (equal), Methodology (equal), Funding acquisition (lead), Supervision (lead), Writing – Review & Editing (equal).

## Data Availability

This study uses data from the Swiss breeding bird survey provided by the Swiss ornithological institute, Sempach. All scripts and data required to run the presented analyses can be accessed from the public GitHub repository https://github.com/UP-macroecology/Malchow_IBMcalibration_2023. Upon acceptance, we will create a permanent Zenodo archive and provide its DOI. The utilised R packages are open-source software: RangeShiftR is available from GitHub. For this study, a tagged development version was used that is available at: BayesianTools is available directly from CRAN or from GitHub under https://github.com/florianhartig/BayesianTools.

## References

1. Aebischer, A. and Scherler, P. (2021). Der Rotmilan - Ein Greifvogel im Aufwind. 1st ed. Bern: Haupt Verlag. 232 pp. isbn: 978-3-258-08249-3.

2. Andrieu, C. and Roberts, G. O. (2009). “The pseudo-marginal approach for efficient Monte Carlo computations”. The Annals of Statistics 37.2, pp. 697–725. doi: 10.1214/07-AOS574.

3. Araújo, M. B. and Peterson, A. T. (2012). “Uses and misuses of bioclimatic envelope modeling”. Ecology 93.7, pp. 1527–1539. doi: 10.1890/11-1930.1.

4. Arlot, S. and Celisse, A. (2010). “A survey of cross-validation procedures for model selection”. Statistics Surveys 4 (none), pp. 40–79. doi: 10.1214/09-SS054.

5. Barber-O’Malley, B., Lassalle, G., Chust, G., Diaz, E., O’Malley, A., Paradinas Blázquez, C., Pórtoles Marquina, J., and Lambert, P. (2022). “HyDiaD: A hybrid species distribution model combining dispersal, multi-habitat suitability, and population dynamics for diadromous species under climate change scenarios”. Ecological Modelling 470, p. 109997. doi: 10.1016/j.ecolmodel.2022.109997.

6. Baucks, C. (2018). “The effect of food supplementation on range use of breeding red kites (Milvus milvus) in Switzerland”. Master thesis. Master thesis. Swiss Ornithological Institute Sempach.

7. Beaumont, M. A. (2010). “Approximate Bayesian Computation in Evolution and Ecology”. Annual Review of Ecology, Evolution, and Systematics 41.1, pp. 379–406. doi: 10.1146/annurev-ecolsys-102209-144621.

8. Bleyhl, B., Ghoddousi, A., Askerov, E., Bocedi, G., Breitenmoser, U., Manvelyan, K., Palmer, S. C. F., Soofi, M., Weinberg, P., Zazanashvili, N., Shmunk, V., Zurell, D., and Kuemmerle, T. (2021). “Reducing persecution is more effective for restoring large carnivores than restoring their prey”. Ecological Applications 31.5, e02338. doi: 10.1002/eap.2338.

9. Bocedi, G., Palmer, S. C. F., Malchow, A.-K., Zurell, D., Watts, K., and Travis, J. M. J. (2021). “RangeShifter 2.0: an extended and enhanced platform for modelling spatial eco-evolutionary dynamics and species’ responses to environmental changes”. Ecography 44.10, pp. 1453–1462. doi: 10.1111/ecog.05687.

10. Bolam, F. C. et al. (2021). “How many bird and mammal extinctions has recent conservation action prevented?” Conservation Letters 14.1, e12762. doi: 10.1111/conl.12762.

11. Braak, C. J. F. ter and Vrugt, J. A. (2008). “Differential Evolution Markov Chain with snooker updater and fewer chains”. Statistics and Computing 18.4, pp. 435–446. doi: 10.1007/s11222-008-9104-9.

12. Briscoe, N. J., Zurell, D., Elith, J., König, C., Fandos, G., Malchow, A.-K., Kéry, M., Schmid, H., and Guillera-Arroita, G. (2021). “Can dynamic occupancy models improve predictions of species’ range dynamics? A test using Swiss birds”. Global Change Biology 27.18, pp. 4269–4282. doi: 10.1111/gcb.15723.

13. Briscoe, N. J. et al. (2019). “Forecasting species range dynamics with process-explicit models: matching methods to applications”. Ecology Letters 22.11, pp. 1940–1956. doi: 10.1111/ele.13348

14. Broms, K. M., Hooten, M. B., Johnson, D. S., Altwegg, R., and Conquest, L. L. (2016). “Dynamic occupancy models for explicit colonization processes”. Ecology 97.1, pp. 194–204. doi: 10.1890/15-0416.1.

15. Cailleret, M., Bircher, N., Hartig, F., Hülsmann, L., and Bugmann, H. (2020). “Bayesian calibration of a growth-dependent tree mortality model to simulate the dynamics of European temperate forests”. Ecological Applications 30.1, e02021. doi: 10.1002/eap.2021.

16. Cereghetti, E., Scherler, P., Fattebert, J., and Grüebler, M. U. (2019). “Quantification of anthropogenic food subsidies to an avian facultative scavenger in urban and rural habitats”. Landscape and Urban Planning 190, p. 103606. doi: 10.1016/j.landurbplan.2019.103606.

17. Csilléry, K., François, O., and Blum, M. G. (2012). “abc: an R package for approximate Bayesian computation (ABC)”. Methods in ecology and evolution 3.3, pp. 475–479.

18. DeAngelis, D. L. and Mooij, W. M. (2005). “Individual-Based Modeling of Ecological and Evolutionary Processes”. Annual Review of Ecology, Evolution, and Systematics 36.1, pp. 147–168. doi: 10.1146/annurev.ecolsys.36.102003.152644.

19. Díaz, S. et al. (2019). “Pervasive human-driven decline of life on Earth points to the need for transformative change”. Science 366.6471, eaax3100. doi: 10.1126/science.aax3100.

20. Dominguez Almela, V., Palmer, S. C. F., Gillingham, P. K., Travis, J. M. J., and Britton, J. R. (2020). “Integrating an individual-based model with approximate Bayesian computation to predict the invasion of a freshwater fish provides insights into dispersal and range expansion dynamics”. Biological Invasions 22.4, pp. 1461–1480. doi: 10.1007/s10530-020-02197-6.

21. Dormann, C. F., Schymanski, S. J., Cabral, J., Chuine, I., Graham, C., Hartig, F., Kearney, M., Morin, X., Römermann, C., Schröder, B., and Singer, A. (2012). “Correlation and process in species distribution models: bridging a dichotomy”. Journal of Biogeography 39.12, pp. 2119–2131. doi: 10.1111/j.1365-2699.2011.02659.x.

22. Dormann, C. F. et al. (2007). “Methods to account for spatial autocorrelation in the analysis of species distributional data: a review”. Ecography 30.5, pp. 609–628. doi: 10.1111/j.2007.0906-7590.05171.x.

23. Dormann, C. F. et al. (2013). “Collinearity: a review of methods to deal with it and a simulation study evaluating their performance”. Ecography 36.1, pp. 27–46. doi: 10.1111/j.1600-0587.2012.07348.x.

24. Duarte, C. M., Agusti, S., Barbier, E., Britten, G. L., Castilla, J. C., Gattuso, J.-P., Fulweiler, R. W., Hughes, T. P., Knowlton, N., Lovelock, C. E., Lotze, H. K., Predragovic, M., Poloczanska, E., Roberts, C., and Worm, B. (2020). “Rebuilding marine life”. Nature 580.7801, pp. 39–51. doi: 10.1038/s41586-020-2146-7.

25. Elith, J. and Leathwick, J. R. (2009). “Species Distribution Models: Ecological Explanation and Prediction Across Space and Time”. Annual Review of Ecology, Evolution, and Systematics 40.1, pp. 677–697. doi: 10.1146/annurev.ecolsys.110308.120159.

26. European Union (2022). Copernicus Land Monitoring Service. European Environment Agency (EEA).

27. Fordham, D. A., Haythorne, S., Brown, S. C., Buettel, J. C., and Brook, B. W. (2021). “poems: R package for simulating species’ range dynamics using pattern-oriented validation”. Methods in Ecology and Evolution 12.12, pp. 2364–2371. doi: 10.1111/2041-210X.13720.

28. Franklin, J. (2013). “Species distribution models in conservation biogeography: developments and challenges”. Diversity and Distributions 19.10, pp. 1217–1223. doi: 10.1111/ddi.12125.

29. Gallien, L., Münkemüller, T., Albert, C. H., Boulangeat, I., and Thuiller, W. (2010). “Predicting potential distributions of invasive species: where to go from here?” Diversity and Distributions 16.3, pp. 331–342. doi: 10.1111/j.1472-4642.2010.00652.x.

30. Gelman, A. and Rubin, D. B. (1992). “Inference from Iterative Simulation Using Multiple Sequences”. Statistical Science 7.4, pp. 457–472. doi: 10.1214/ss/1177011136.

31. Grimm, V. et al. (2020). “The ODD protocol for describing agent-based and other simulation models: A second update to improve clarity, replication, and structural realism”. Journal of Artificial Societies and Social Simulation 23.2.

32. Grimm, V., Revilla, E., Berger, U., Jeltsch, F., Mooij, W. M., Railsback, S. F., Thulke, H.-H., Weiner, J., Wiegand, T., and DeAngelis, D. L. (2005). “Pattern-Oriented Modeling of Agent-Based Complex Systems: Lessons from Ecology”. Science 310.5750, pp. 987–991. doi: 10.1126/science.1116681.

33. Guisan, A. and Thuiller, W. (2005). “Predicting species distribution: offering more than simple habitat models”. Ecology Letters 8.9, pp. 993–1009. doi: 10.1111/j.1461-0248.2005.00792.x.

34. Guisan, A. et al. (2013). “Predicting species distributions for conservation decisions”. Ecology Letters 16.12. Ed. by H. Arita, pp. 1424–1435. doi: 10.1111/ele.12189.

35. Hagen, O., Flück, B., Fopp, F., Cabral, J. S., Hartig, F., Pontarp, M., Rangel, T. F., and Pellissier, L. (2021). “gen3sis: A general engine for eco-evolutionary simulations of the processes that shape Earth’s biodiversity”. PLOS Biology 19.7, e3001340. doi: 10.1371/journal.pbio.3001340.

36. Hartig, F., Dyke, J., Hickler, T., Higgins, S. I., O’Hara, R. B., Scheiter, S., and Huth, A. (2012). “Connecting dynamic vegetation models to data – an inverse perspective”. Journal of Biogeography 39.12, pp. 2240–2252. doi: 10.1111/j.1365-2699.2012.02745.x.

37. Hartig, F., Minunno, F., and Paul, S. (2019). “BayesianTools: General-purpose MCMC and SMC Samplers and Tools for Bayesian Statistics”. R package version 0.1.7.

38. Hauenstein, S., Fattebert, J., Grüebler, M. U., Naef-Daenzer, B., Pe’er, G., and Hartig, F. (2019). “Calibrating an individual-based movement model to predict functional connectivity for little owls”. Ecological Applications 29.4, e01873. doi: 10.1002/eap.1873.

39. Hefley, T. J., Hooten, M. B., Russell, R. E., Walsh, D. P., and Powell, J. A. (2017). “When mechanism matters: Bayesian forecasting using models of ecological diffusion”. Ecology Letters 20.5, pp. 640–650.

40. Hobbs, N. T. and Hooten, M. B. (2015). Bayesian models: a statistical primer for ecologists. Princeton University Press.

41. Hoffmann, M. et al. (2010). “The Impact of Conservation on the Status of the World’s Vertebrates”. Science 330.6010, pp. 1503–1509. doi: 10.1126/science.1194442.

42. Jaatinen, K., Westerbom, M., Norkko, A., Mustonen, O., and Koons, D. N. (2021). “Detrimental impacts of climate change may be exacerbated by density-dependent population regulation in blue mussels”. Journal of Animal Ecology 90.3, pp. 562–573. doi: 10.1111/1365-2656.13377.

43. Karger, D. N., Conrad, O., Böhner, J., Kawohl, T., Kreft, H., Soria-Auza, R. W., Zimmermann, N. E., Linder, H. P., and Kessler, M. (2017). “Climatologies at high resolution for the earth’s land surface areas”. Scientific Data 4, p. 170122. doi: 10.1038/sdata.2017.122.

44. Karger, D. N., Conrad, O., Böhner, J., Kawohl, T., Kreft, H., Soria-Auza, R. W., Zimmermann, N. E., Linder, H. P., and Kessler, M. (2018). Data from: Climatologies at high resolution for the earth’s land surface areas. doi: 10.5061/dryad.kd1d4.

45. Katzenberger, J., Gottschalk, E., Balkenhol, N., and Waltert, M. (2019). “Long-term decline of juvenile survival in German Red Kites”. Journal of Ornithology 160.2, pp. 337–349. doi: 10.1007/s10336-018-1619-z.

46. Kearney, M. and Porter, W. (2009). “Mechanistic niche modelling: combining physiological and spatial data to predict species’ ranges”. Ecology Letters 12.4, pp. 334–350. doi: 10.1111/j.1461-0248.2008.01277.x.

47. Keith, D. A., Akçakaya, H. R., Thuiller, W., Midgley, G. F., Pearson, R. G., Phillips, S. J., Regan, H. M., Araújo, M. B., and Rebelo, T. G. (2008). “Predicting extinction risks under climate change: coupling stochastic population models with dynamic bioclimatic habitat models”. Biology Letters 4.5, pp. 560–563. doi: 10.1098/rsbl.2008.0049.

48. Kéry, M., Guillera-Arroita, G., and Lahoz-Monfort, J. J. (2013). “Analysing and mapping species range dynamics using occupancy models”. Journal of Biogeography 40.8, pp. 1463–1474. doi: 10.1111/jbi.12087.

49. Kissling, W. D. et al. (2018). “Building essential biodiversity variables (EBVs) of species distribution and abundance at a global scale”. Biological Reviews 93.1, pp. 600–625. doi: 10.1111/brv.12359.

50. Knaus, P., Strebel, N., and Sattler, T. (2022). The State of Birds in Switzerland 2022. Sempach: Swiss Ornithological Institute.

51. Knaus, P., Jérôme, G., Sattler, T., Wechsler, S., Kéry, M., Strebel, N., and Antoniazza, S. (2018). Schweizer Brutvogelatlas 2013-2016. Sempach, Schweiz: Schweizerische Vogelwarte. isbn: 978-3-85949-009-3.

52. Landguth, E. L., Bearlin, A., Day, C. C., and Dunham, J. (2017). “CDMetaPOP: an individual-based, eco-evolutionary model for spatially explicit simulation of landscape demogenetics”. Methods in Ecology and Evolution 8.1, pp. 4–11. doi: 10.1111/2041-210X.12608.

53. Luengo, D., Martino, L., Bugallo, M., Elvira, V., and Särkkä, S. (2020). “A survey of Monte Carlo methods for parameter estimation”. EURASIP Journal on Advances in Signal Processing 2020.1, p. 25. doi: 10.1186/s13634-020-00675-6.

54. Malchow, A.-K., Bocedi, G., Palmer, S. C. F., Travis, J. M. J., and Zurell, D. (2021). “RangeShiftR: an R package for individual-based simulation of spatial eco-evolutionary dynamics and species’ responses to environmental changes”. Ecography 44.10, pp. 1443–1452. doi: 10.1111/ecog.05689.

55. Malchow, A.-K., Hartig, F., Reeg, J., Kéry, M., and Zurell, D. (2023). “Demography-environment relationships improve mechanistic understanding of range dynamics under climate change”. Philosophical Transactions of the Royal Society B. doi: 10.1098/rstb.2022.0194.

56. Marion, G., McInerny, G. J., Pagel, J., Catterall, S., Cook, A. R., Hartig, F., and O’Hara, R. B. (2012). “Parameter and uncertainty estimation for process-oriented population and distribution models: data, statistics and the niche”. Journal of Biogeography 39.12, pp. 2225–2239. doi: 10.1111/j.1365-2699.2012.02772.x.

57. Morris, M. D. (1991). “Factorial Sampling Plans for Preliminary Computational Experiments”. Technometrics 33.2, pp. 161–174. doi: 10.1080/00401706.1991.10484804.

58. Mortensen, L. O., Chudzinska, M. E., Slabbekoorn, H., and Thomsen, F. (2021). “Agent-based models to investigate sound impact on marine animals: bridging the gap between effects on individual behaviour and population level consequences”. Oikos 130.7, pp. 1074–1086. doi: 10.1111/oik.08078.

59. Moulin, T., Perasso, A., Calanca, P., and Gillet, F. (2021). “DynaGraM: A process-based model to simulate multi-species plant community dynamics in managed grasslands”. Ecological Modelling 439, p. 109345. doi: 10.1016/j.ecolmodel.2020.109345.

60. Nachtigall, W. (2008). “Der Rotmilan (Milvus milvus, L. 1758) in Sachsen und Südbrandenburg–Untersuchungen zu Verbreitung und Ökologie”. Diss., Univ. Halle-Wittenberg.

61. Nägeli, M., Scherler, P., Witczak, S., Catitti, B., Aebischer, A., Bergen, V. van, Kormann, U., and Grüebler, M. U. (2021). “Weather and food availability additively affect reproductive output in an expanding raptor population”. Oecologia. doi: 10.1007/s00442-021-05076-6.

62. Newbold, T. et al. (2015). “Global effects of land use on local terrestrial biodiversity”. Nature 520.7545, pp. 45–50. doi: 10.1038/nature14324.

63. Newson, R. (2006). “Confidence Intervals for Rank Statistics: Somers’ D and Extensions”. The Stata Journal 6.3, pp. 309–334. doi: 10.1177/1536867X0600600302.

64. Newton, I., Davis, P., and Davis, J. (1989). “Age of first breeding, dispersal and survival of Red Kites Milvus milvus in Wales”. Ibis 131.1, pp. 16–21.

65. Oliver, T. H., Gillings, S., Girardello, M., Rapacciuolo, G., Brereton, T. M., Siriwardena, G. M., Roy, D. B., Pywell, R., and Fuller, R. J. (2012). “Population density but not stability can be predicted from species distribution models”. Journal of Applied Ecology 49.3, pp. 581–590. doi: 10.1111/j.1365-2664.2012.02138.x.

66. Pellissier, L., Rohr, R. P., Ndiribe, C., Pradervand, J.-N., Salamin, N., Guisan, A., and Wisz, M. (2013). “Combining food web and species distribution models for improved community projections”. Ecology and Evolution 3.13, pp. 4572–4583. doi: 10.1002/ece3.843.

67. Pfeiffer, T. and Schaub, M. (2023). “Productivity drives the dynamics of a red kite source population that depends on immigration”. Journal of Avian Biology 2023.1–2, e02984.

68. R Core Team (2020). R: A Language and Environment for Statistical Computing. Vienna, Austria: R Foundation for Statistical Computing.

69. Railsback, S. F. and Grimm, V. (2019). Agent-Based and Individual-Based Modeling: A Practical Introduction, Second Edition. Princeton University Press. 359 pp. isbn: 978-0-691-19004-4.

70. Risk, B. B., Valpine, P. de, and Beissinger, S. R. (2011). “A robust-design formulation of the incidence function model of metapopulation dynamics applied to two species of rails”. Ecology 92.2, pp. 462–474. doi: 10.1890/09-2402.1.

71. Roberts, D. R., Bahn, V., Ciuti, S., Boyce, M. S., Elith, J., Guillera-Arroita, G., Hauenstein, S., Lahoz-Monfort, J. J., Schröder, B., Thuiller, W., Warton, D. I., Wintle, B. A., Hartig, F., and Dormann, C. F. (2017). “Cross-validation strategies for data with temporal, spatial, hierarchical, or phylogenetic structure”. Ecography 40.8, pp. 913–929. doi: 10.1111/ecog.02881.

72. Rodríguez, L., García, J. J., Carreño, F., and Martínez, B. (2019). “Integration of physiological knowledge into hybrid species distribution modelling to improve forecast of distributional shifts of tropical corals”. Diversity and Distributions 25.5, pp. 715–728. doi: 10.1111/ddi.12883.

73. Santos, E. P., Wagner, H. H., Ferraz, S. F. B., and Siqueira, T. (2020). “Interactive persistent effects of past land-cover and its trajectory on tropical freshwater biodiversity”. Journal of Applied Ecology 57.11, pp. 2149–2158. doi: 10.1111/1365-2664.13717.

74. Schaub, M. (2012). “Spatial distribution of wind turbines is crucial for the survival of red kite populations”. Biological Conservation 155, pp. 111–118. doi: 10.1016/j.biocon.2012.06.021.

75. Schmid, H., Luder, R., Naef-Daenzer, B., Graf, R., and Zbinden, N. (1998). Schweizer Brutvogelatlas 1993–1996. Sempach, Schweiz: Schweizerische Vogelwarte. isbn: 978-3-9521064-5-7.

76. Schmid, H., Zbinden, N., and Keller, V. (2004). Überwachung der Bestandsentwicklung häufiger Brutvögel in der Schweiz. Sempach: Schweizerische Vogelwarte.

77. Schmolke, A., Thorbek, P., DeAngelis, D. L., and Grimm, V. (2010). “Ecological models supporting environmental decision making: a strategy for the future”. Trends in Ecology & Evolution 25.8, pp. 479–486. doi: 10.1016/j.tree.2010.05.001.

78. Schurr, F. M., Pagel, J., Cabral, J. S., Groeneveld, J., Bykova, O., O’Hara, R. B., Hartig, F., Kissling, W. D., Linder, H. P., Midgley, G. F., Schröder, B., Singer, A., and Zimmermann, N. E. (2012). “How to understand species’ niches and range dynamics: a demographic research agenda for biogeography”. Journal of Biogeography 39.12, pp. 2146–2162. doi: 10.1111/j.1365-2699.2012.02737.x.

79. Schweiger, O., Heikkinen, R. K., Harpke, A., Hickler, T., Klotz, S., Kudrna, O., Kühn, I., Pöyry, J., and Settele, J. (2012). “Increasing range mismatching of interacting species under global change is related to their ecological characteristics”. Global Ecology and Biogeography 21.1, pp. 88–99. doi: 10.1111/j.1466-8238.2010.00607.x.

80. Selwood, K. E., McGeoch, M. A., and Mac Nally, R. (2015). “The effects of climate change and land-use change on demographic rates and population viability”. Biological Reviews 90.3, pp. 837–853. doi: 10.1111/brv.12136.

81. Semper-Pascual, A., Burton, C., Baumann, M., Decarre, J., Gavier-Pizarro, G., Gómez-Valencia, B., Macchi, L., Mastrangelo, M. E., Pötzschner, F., Zelaya, P. V., and Kuemmerle, T. (2021). “How do habitat amount and habitat fragmentation drive time-delayed responses of biodiversity to land-use change?” Proceedings of the Royal Society B: Biological Sciences 288.1942, p. 20202466. doi: 10.1098/rspb.2020.2466.

82. Sergio, F., Tavecchia, G., Blas, J., Tanferna, A., and Hiraldo, F. (2021). “Demographic modeling to fine-tune conservation targets: importance of pre-adults for the decline of an endangered raptor”. Ecological Applications 31.3, e2266. doi: 10.1002/eap.2266.

83. Sillero, N., Arenas-Castro, S., Enriquez-Urzelai, U., Vale, C. G., Sousa-Guedes, D., Martínez-Freiría, F., Real, R., and Barbosa, A. M. (2021). “Want to model a species niche? A step-by-step guideline on correlative ecological niche modelling”. Ecological Modelling 456, p. 109671. doi: 10.1016/j.ecolmodel.2021.109671.

84. Singer, A., Schweiger, O., Kühn, I., and Johst, K. (2018). “Constructing a hybrid species distribution model from standard large-scale distribution data”. Ecological Modelling 373, pp. 39–52. doi: 10.1016/j.ecolmodel.2018.02.002.

85. Smolik, M., Dullinger, S., Essl, F., Kleinbauer, I., Leitner, M., Peterseil, J., Stadler, L.-M., and Vogl, G. (2010). “Integrating species distribution models and interacting particle systems to predict the spread of an invasive alien plant”. Journal of Biogeography 37.3, pp. 411–422. doi: 10.1111/j.1365-2699.2009.02227.x.

86. Thompson, B. K., Olden, J. D., and Converse, S. J. (2021). “Mechanistic invasive species management models and their application in conservation”. Conservation Science and Practice 3.11, e533. doi: 10.1111/csp2.533.

87. Travis, J. M. J., Mustin, K., Bartoń, K. A., Benton, T. G., Clobert, J., Delgado, M. M., Dytham, C., Hovestadt, T., Palmer, S. C. F., Dyck, H. V., and Bonte, D. (2012). “Modelling dispersal: an eco-evolutionary framework incorporating emigration, movement, settlement behaviour and the multiple costs involved”. Methods in Ecology and Evolution 3.4, pp. 628–641. doi: 10.1111/j.2041-210X.2012.00193.x.

88. Urban, M. C. et al. (2016). “Improving the forecast for biodiversity under climate change”. Science 353.6304. doi: 10.1126/science.aad8466.

89. Valavi, R., Elith, J., Lahoz-Monfort, J. J., and Guillera-Arroita, G. (2019). “blockCV: An r package for generating spatially or environmentally separated folds for k-fold cross-validation of species distribution models”. Methods in Ecology and Evolution 10.2, pp. 225–232. doi: 10.1111/2041-210X.13107.

90. Vehtari, A., Gelman, A., and Gabry, J. (2017). “Practical Bayesian model evaluation using leave-one-out cross-validation and WAIC”. Statistics and Computing 27.5, pp. 1413–1432. doi: 10.1007/s11222-016-9696-4.

91. Vehtari, A., Gelman, A., Simpson, D., Carpenter, B., and Bürkner, P.-C. (2021). “Rank-Normalization, Folding, and Localization: An Improved R^ for Assessing Convergence of MCMC (with Discussion)”. Bayesian Analysis 16.2, pp. 667–718. doi: 10.1214/20-BA1221.

92. Visintin, C., Briscoe, N. J., Woolley, S. N. C., Lentini, P. E., Tingley, R., Wintle, B. A., and Golding, N. (2020). “steps: Software for spatially and temporally explicit population simulations”. Methods in Ecology and Evolution 11.4, pp. 596–603. doi: 10.1111/2041-210X.13354.

93. Watts, K., Whytock, R. C., Park, K. J., Fuentes-Montemayor, E., Macgregor, N. A., Duffield, S., and McGowan, P. J. K. (2020). “Ecological time lags and the journey towards conservation success”. Nature Ecology & Evolution 4.3, pp. 304–311. doi: 10.1038/s41559-019-1087-8.

94. Wenger, S. J. and Olden, J. D. (2012). “Assessing transferability of ecological models: an underappreciated aspect of statistical validation”. Methods in Ecology and Evolution 3.2, pp. 260–267. doi: 10.1111/j.2041-210X.2011.00170.x.

95. Wikle, C. K. (2003). “Hierarchical Bayesian models for predicting the spread of ecological processes”. Ecology 84.6, pp. 1382–1394.

96. Zurell, D., König, C., Malchow, A.-K., Kapitza, S., Bocedi, G., Travis, J., and Fandos, G. (2022). “Spatially explicit models for decision-making in animal conservation and restoration”. Ecography 2022.4. doi: 10.1111/ecog.05787.

97. Zylstra, E. R. and Zipkin, E. F. (2021). “Accounting for sources of uncertainty when forecasting population responses to climate change”. Journal of Animal Ecology 90.3, pp. 558–561. doi: 10.1111/1365-2656.13443.

