## Supplemental Tables and Figures for "Fitting individual-based models of spatial population dynamics to long-term monitoring data"

Ecological Applications – Article

#### S1 Appendix 1: Supplemental Figures and Tables

Table S1: Overview of all data sources used to parameterise and calibrate the dynamic SDM.

| Usage | Data type | Data source |
|---|---|---|
| **Direct parametrisation** | | |
| Initial distribution | Abundance snapshot | Breeding bird Atlas II (1993-96) (Schmid et al., 1998) |
| Static habitat model (cSDM) | Presence/absence | Breeding bird Atlas III (2013-16) (Knaus et al., 2018) |
| | Bioclimatic variables | CHELSA Bioclim v1.2: WorldClim (1979-2013) (Karger et al., 2017; Karger et al., 2018) |
| | Land cover | CORINE (2012) (European Union, 2022) |
| Habitat suitability maps (cSDM projections) | Bioclimatic variables - same as for fitting | CHELSA Bioclim v1.2: WorldClim (1979-2013) |
| | Land cover - varying | CORINE (2000, 2006, 2012, 2018) (European Union, 2022) |
| Individual-based model | Stage structure | Sergio et al. (2021) and Newton et al. (1989) |
| | Survival probabilities | Katzenberger et al. (2019) (also Schaub (2012) and Newton et al. (1989)) |
| | Fecundity | Nägeli et al. (2021) (also Schaub (2012) and Nachtigall (2008)) |
| | Emigration probability | own unpublished data |
| | Dispersal distance | Nachtigall (2008) (also Newton et al. (1989) and own unpublished data) |
| **Indirect parametrisation** | | |
| Calibration (likelihood input) | Abundance time series | Swiss breeding bird survey (1999-2019) (Schmid & Volet, 2004) |
| **Validation** | | |
| Internal validation (cross-validation) | Abundance time series (folds of calibration data set) | Swiss breeding bird survey (1999-2019) (Schmid & Volet, 2004) |
| External validation | Relative abundance time series | Swiss breeding bird index (1999-2019) (Knaus et al., 2022) |

Table S2: The CORINE land cover classes (column 3) were aggregated to the land-use variables (column 2) that were considered as predictors in the habitat suitability model (HSM). The asterisks in column 1 mark those aggregated variables that remained after the variable selection process and were used as predictors in the HSM.

| HSM var | Aggregated land cover type | Corine classes |
|---|---|---|
| | artificial surfaces | 111-142 |
| | arable land | 211-213 |
| * | orchards, pastures, heterogeneous agricultural areas | 221-223, 231, 241-244 |
| * | forests | 311-313 |
| | natural grassland, moors and heathland | 321-324 |
| | no vegetation | 331-335 |
| * | wetlands, water bodies | 411-423, 511-523 |

Table S3: All nineteen WorldClim Bioclimatic variables (column 2 and 3) were considered as predictors in the habitat suitability model (HSM). The asterisks in column 1 mark those variables that remained after the variable selection process and were used as predictors in the HSM.

| HSM var | Name | Description |
|---|---|---|
| | BIO1 | Annual Mean Temperature |
| * | BIO2 | Mean Diurnal Range (monthly Mean($max_{temp}$ - $min_{temp}$)) |
| * | BIO3 | Isothermality (BIO2/BIO7) (* 100) |
| * | BIO4 | Temperature Seasonality (standard deviation *100) |
| * | BIO5 | Max Temperature of Warmest Month |
| | BIO6 | Min Temperature of Coldest Month |
| | BIO7 | Temperature Annual Range |
| | BIO8 | Mean Temperature of Wettest Quarter |
| * | BIO9 | Mean Temperature of Driest Quarter |
| | BIO10 | Mean Temperature of Warmest Quarter |
| | BIO11 | Mean Temperature of Coldest Quarter |
| | BIO12 | Annual Precipitation |
| * | BIO13 | Precipitation of Wettest Month |
| * | BIO14 | Precipitation of Driest Month |
| * | BIO15 | Precipitation Seasonality (Coefficient of Variation) |
| | BIO16 | Precipitation of Wettest Quarter |
| | BIO17 | Precipitation of Driest Quarter |
| | BIO18 | Precipitation of Warmest Quarter |
| | BIO19 | Precipitation of Coldest Quarter |

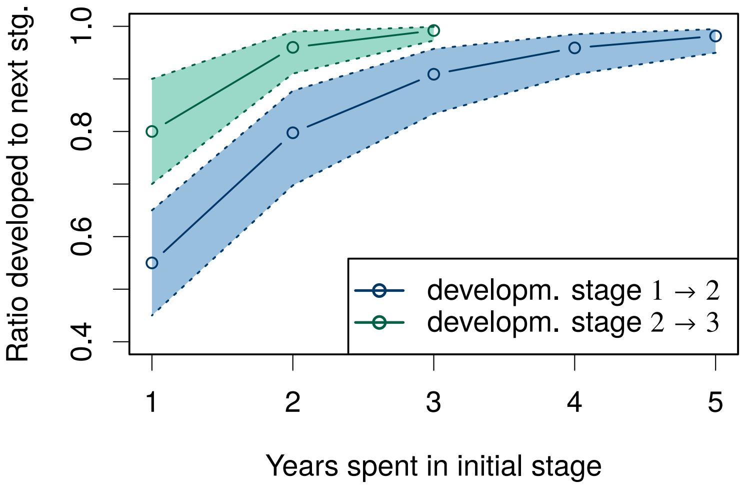


Figure S1: Ratio of individuals of an initial stage (1 or 2) that have developed to the next stage (2 or 3) after a number of years $x$ have passed.

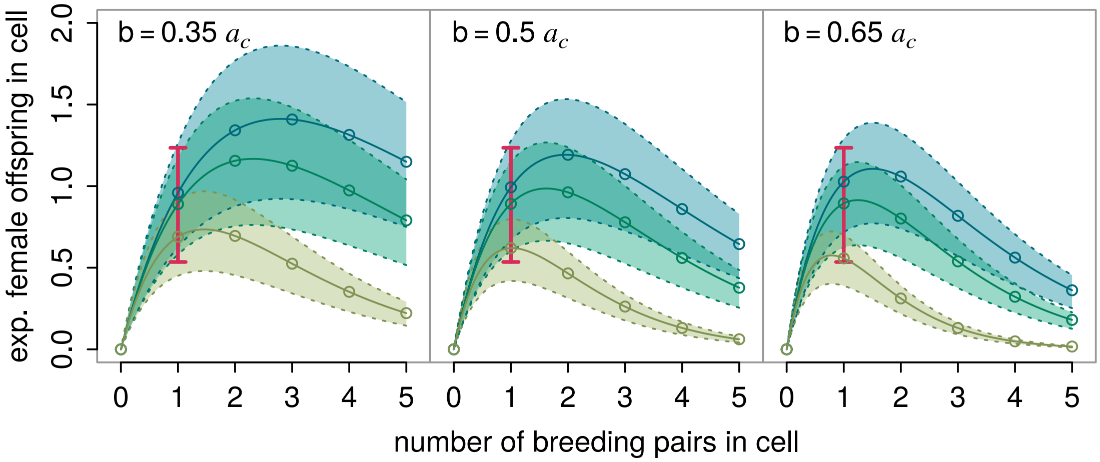


Figure S2: Mean and standard deviation of the number of expected female offspring per cell over the number of breeding pairs in that cell for different values of the parameter that controls the strength of demographic density-dependence $b = (0.50 \pm 0.15)\,a_c$. Blue, turquoise and green show this relation for a cell of 98, 81, and 51% habitat suitability (the maximum, median and minimum HSI over all habitat cells). The red bar indicates the range of observed fecundities $\phi_1 = 0.88 \pm 0.35$.

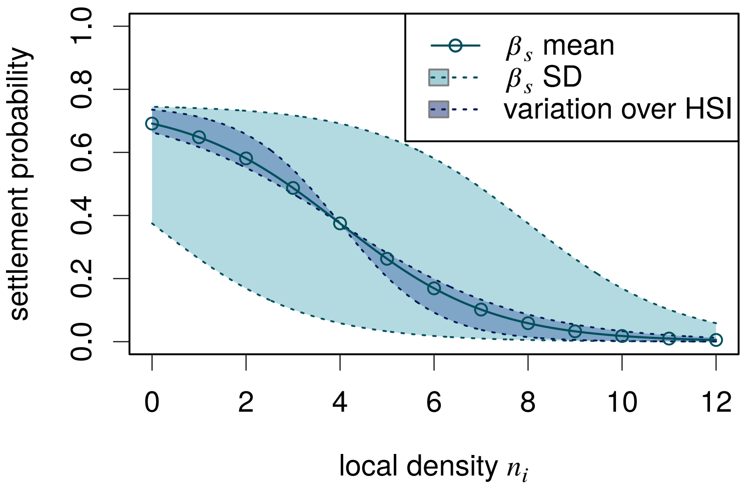


Figure S3: Mean and standard deviation of settlement probability $p_s = p_{s,0}\,(1 + e^{-\alpha_s\,(b_i n_i - \beta_s)})^{-1}$ over the local population density $n_i$ given in breeding pairs per cell. The inflection point $\beta_s$ is a calibrated parameter and its prior estimate is $\hat{\beta}_s = \beta_s/b = (4 \pm 4)\,a_c^{-1}$. The maximum settlement probability $s_0 = 0.75$ and slope parameter $\alpha_s = -1$ are fixed.

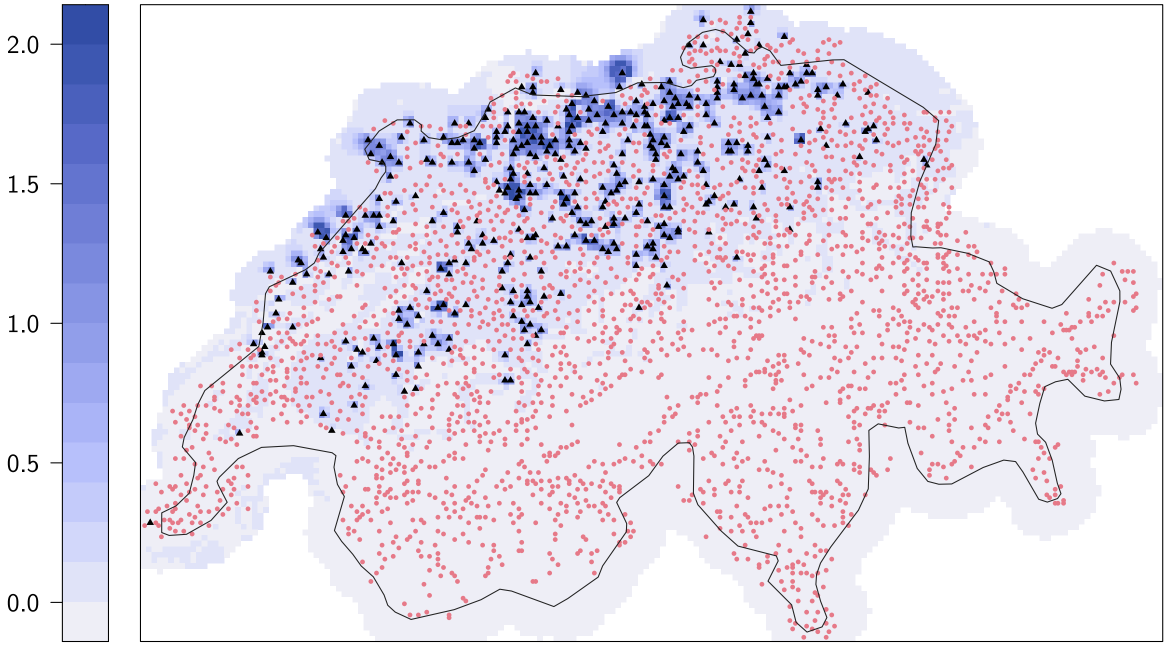


Figure S4: The initial distribution of stage-3 individuals in each simulation is drawn from a joint Poisson distribution. The map shows the respective means for each cell on the colour scale. The underlying Poisson model is based on the Atlas data from 1993-’96; their absences and presences are shown as red circles and black triangles, respectively.

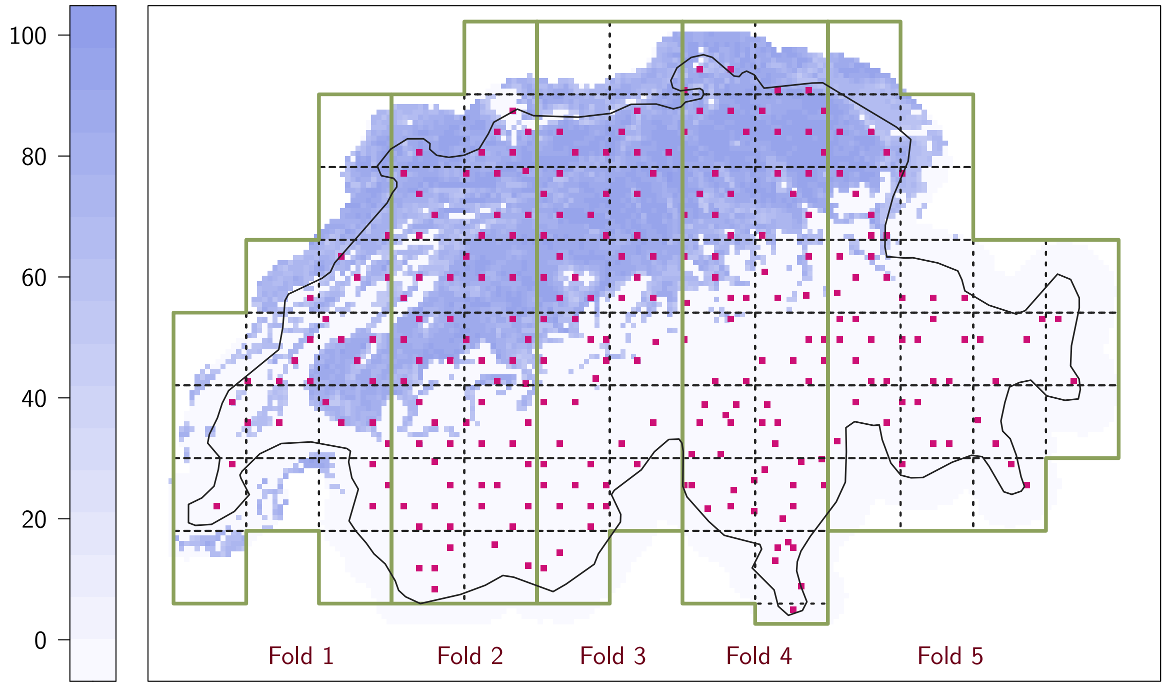


Figure S5: Spatial aggregation of observed and simulated abundance data: The solid green line delineates the spatial folds used for model evaluation. The broken blue line shows the spatial blocks within which the abundance data are aggregated. The map shows the simulation grid with the habitat suitability in year 2018 of each 2×2 $km^2$ cell (blue colour scale) and the locations of MHB sites (pink squares) for reference.

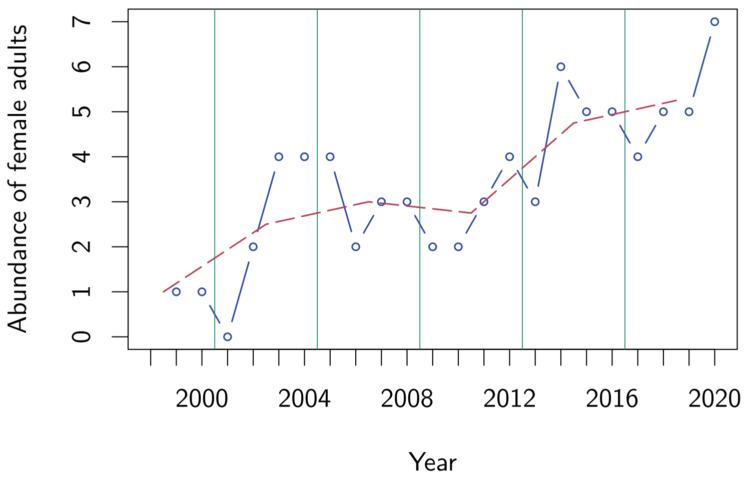


Figure S6: Temporal aggregation of observed and simulated abundance data: The vertical lines delineate the temporal blocks within which the abundance data are aggregated. The blue line and circles show an exemplary time series of total female abundance within a given spatial block, the broken red line shows their means within each temporal block.

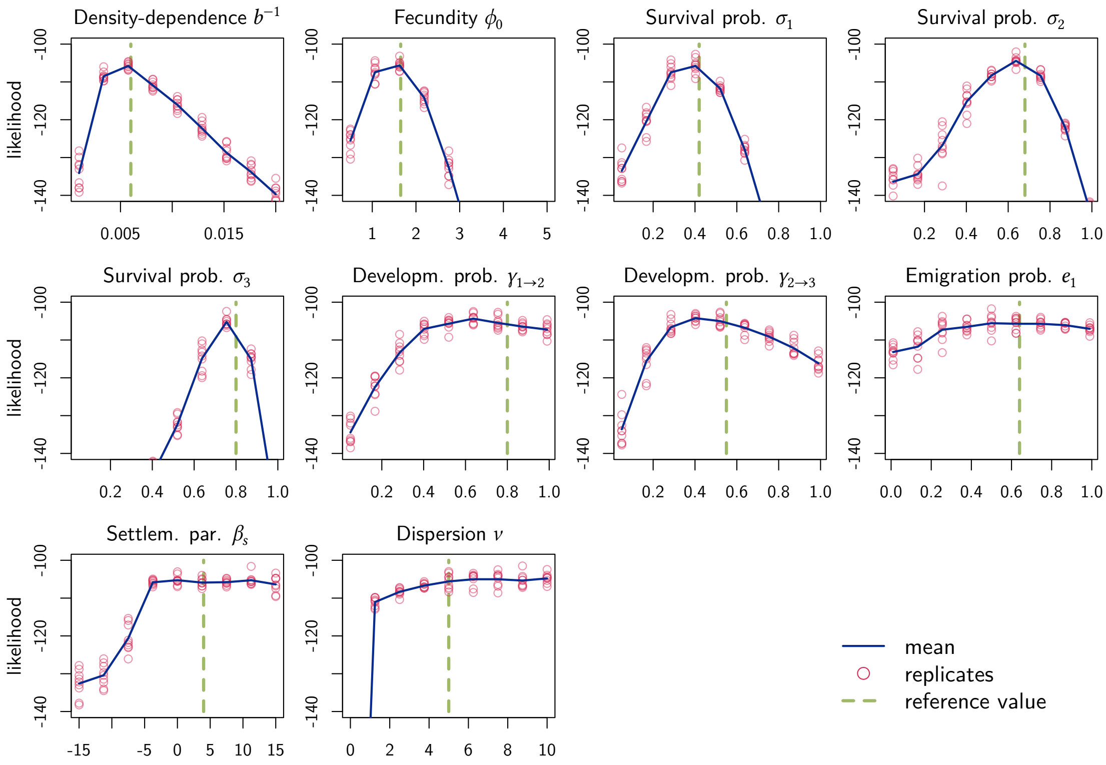


Figure S7: One-factor-at-a-time sensitivity analysis. The green broken vertical lines mark the reference values, with which the reference data $D_{\text{ref}}$ was simulated. To evaluate the sensitivity of each single parameter $\theta_i$ on the likelihood $l(\theta) = p(D_{\text{sim}} \,|\, \theta, M)$, it is varied within the boundaries of its prior while keeping all other parameters constant and its likelihood with respect to $D_{\text{ref}}$ is evaluated.

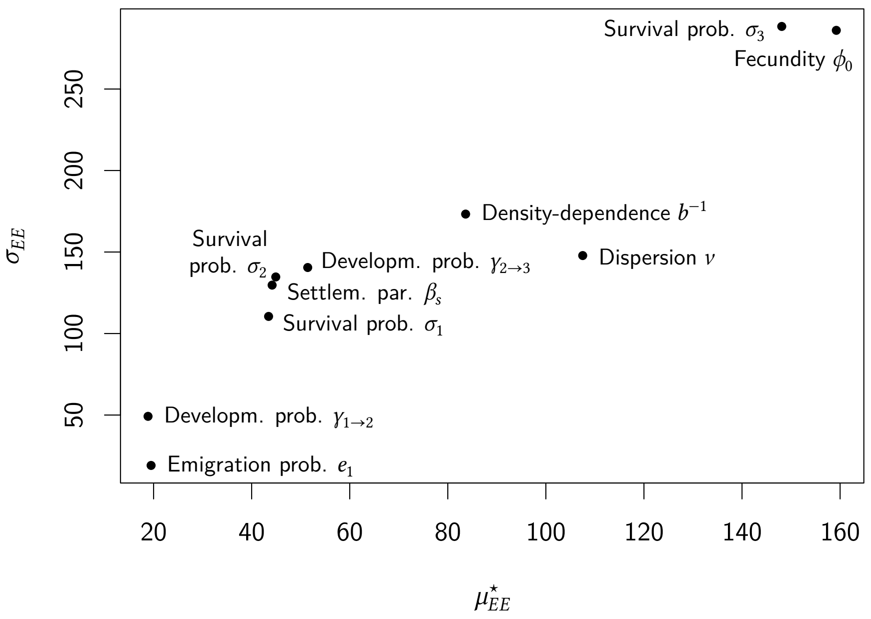


Figure S8: Global sensitivity analysis using Morris' elementary effects screening method (Morris, 1991). Shown are the mean of the absolute value of the elementary effect $\mu^{\star}_{EE}$ and the standard deviation of the elementary effect $\sigma_{EE}$ for each calibrated parameter $\theta_i$. An estimate of the overall sensitivity of $\theta_i$ on the likelihood $l(\theta)$ is provided by $\mu^{\star}_{EE}$, whereas $\sigma_{EE}$ indicates a non-linear relation or potential interactions.

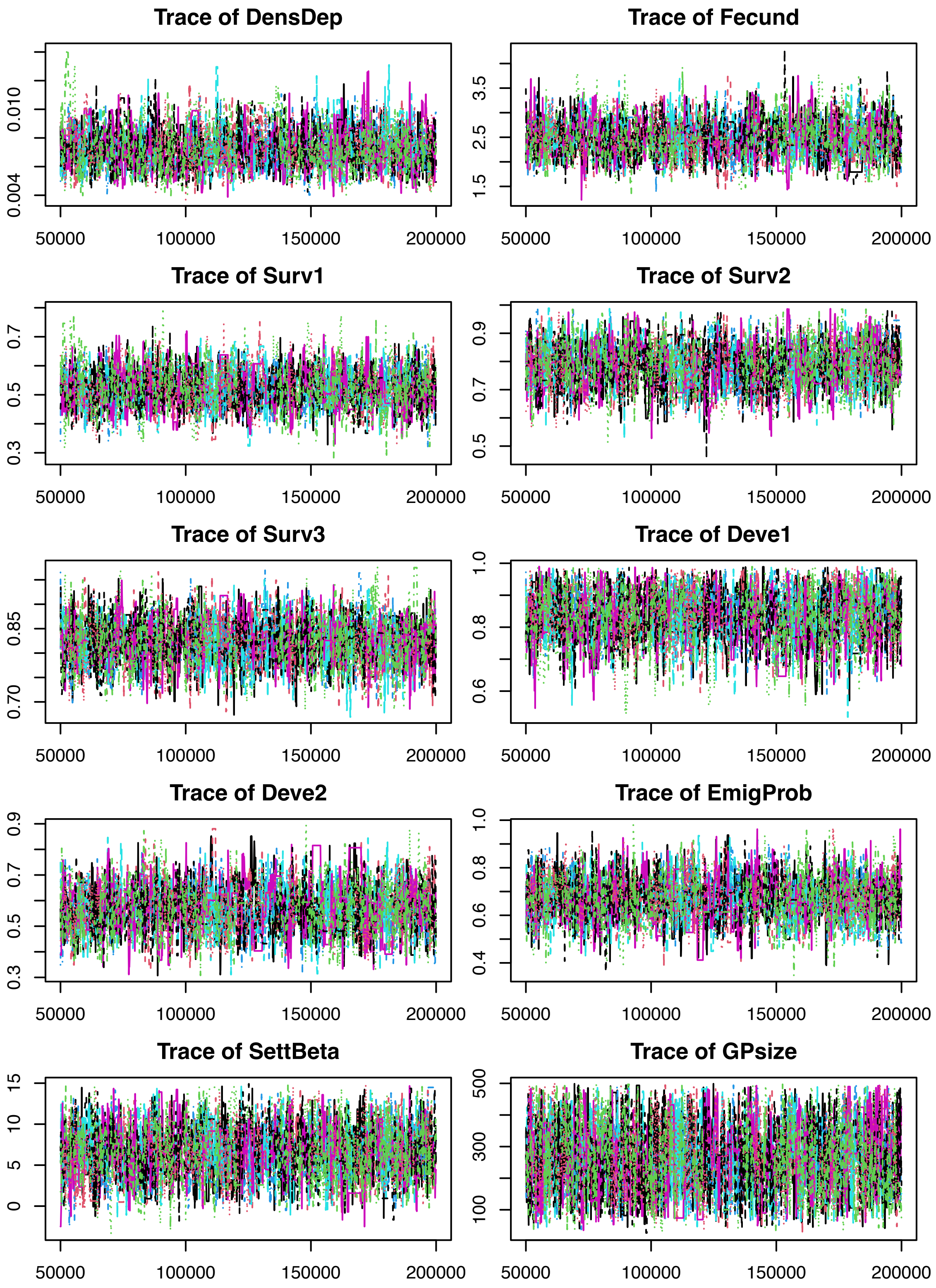


Figure S9: Trace plots: Parameter value sampled by each chain in each iteration.

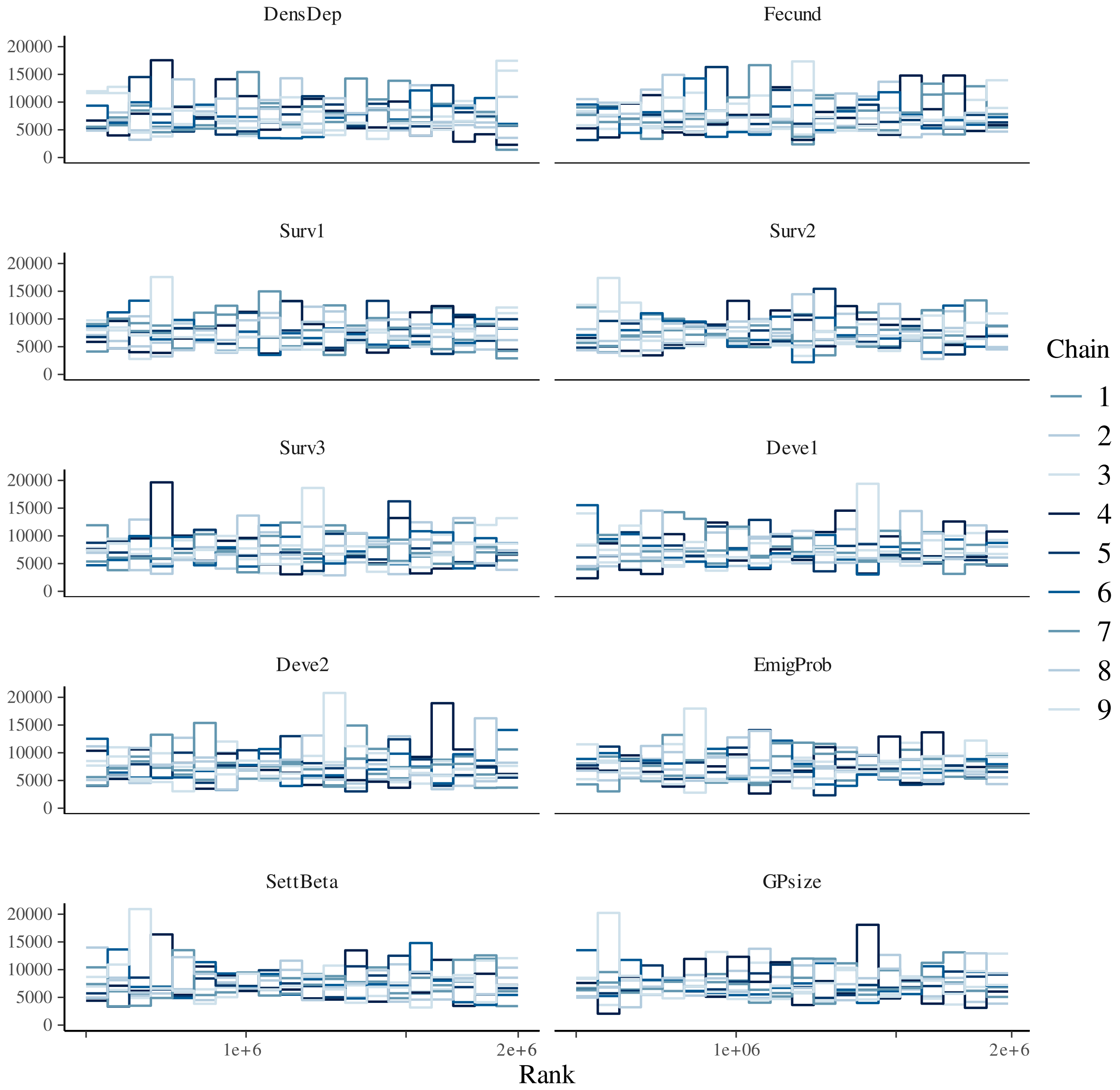


Figure S10: Trace rank plots: Histograms of the rank of the parameter values sampled by each chain, where ranking is done over all draws by all chains (Vehtari et al., 2021).

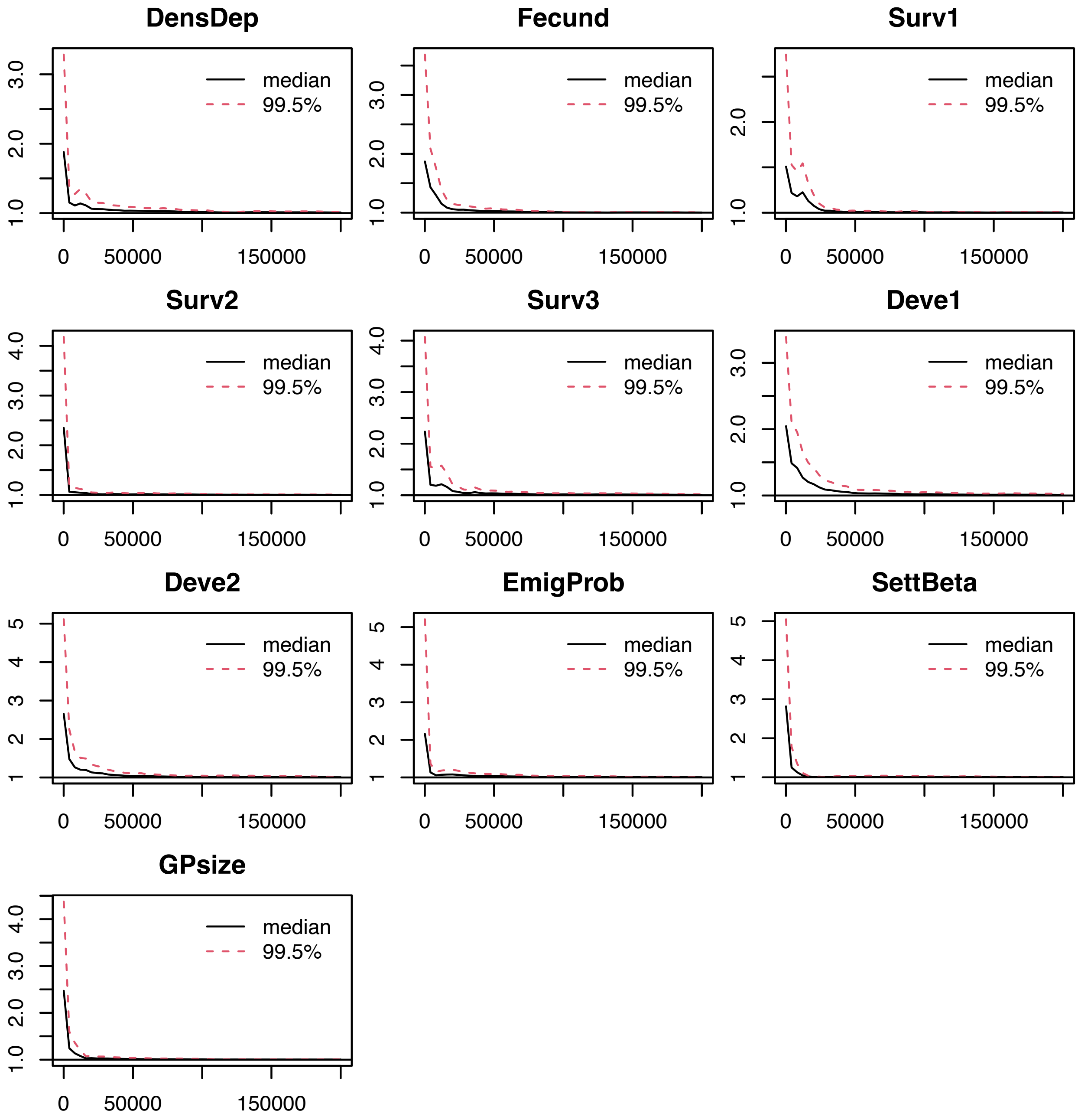


Figure S11: Potential scale reduction factor (psrf; Gelman & Rubin, 1992) at different chain lengths for all calibration parameters. Chain length is $2 \times 10^5$ iterations of which $0.5 \times 10^5$ are discarded as burn-in. Multivariate psrf evaluates to 1.05, 1.02, 1.05, 1.09, and 1.04 for the folds 1 to 5, and to 1.03 for the calibration run using the whole data set.

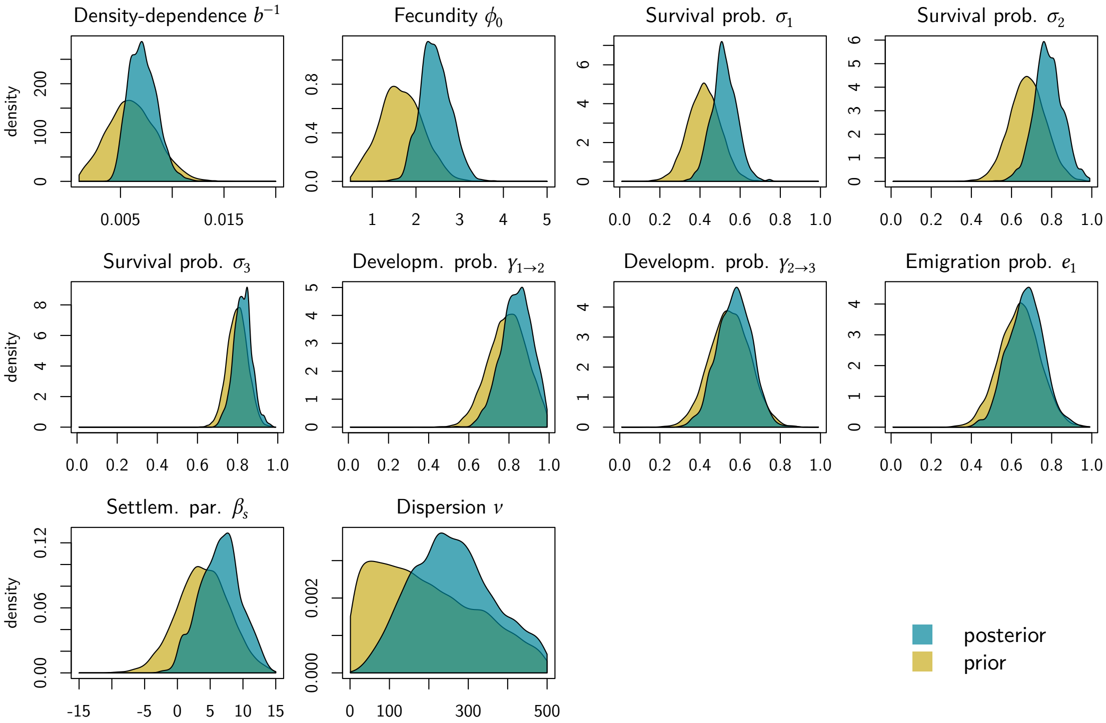


Figure S12: Marginal posterior distributions, fitted to full MHB data set (all five spatial folds). Combination of three independent DEzs chains with three internal chains each.

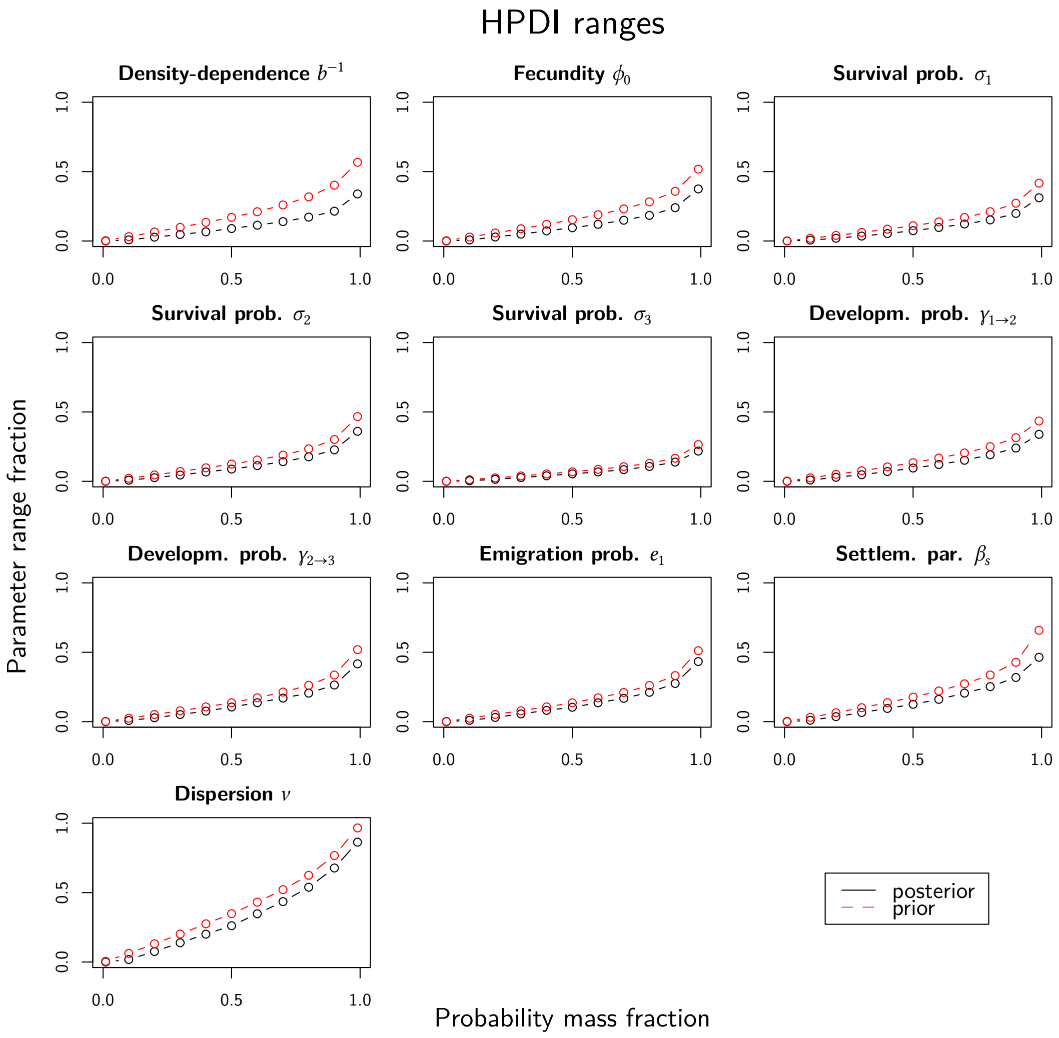


Figure S13: Comparison of the width of posterior versus prior marginals using highest-posterior-density intervals (HPDIs): The smaller the HPDI width and the shallower its slope with increasing probability mass, the tighter is the distribution, translating to less uncertainty around the parameters' point estimate.

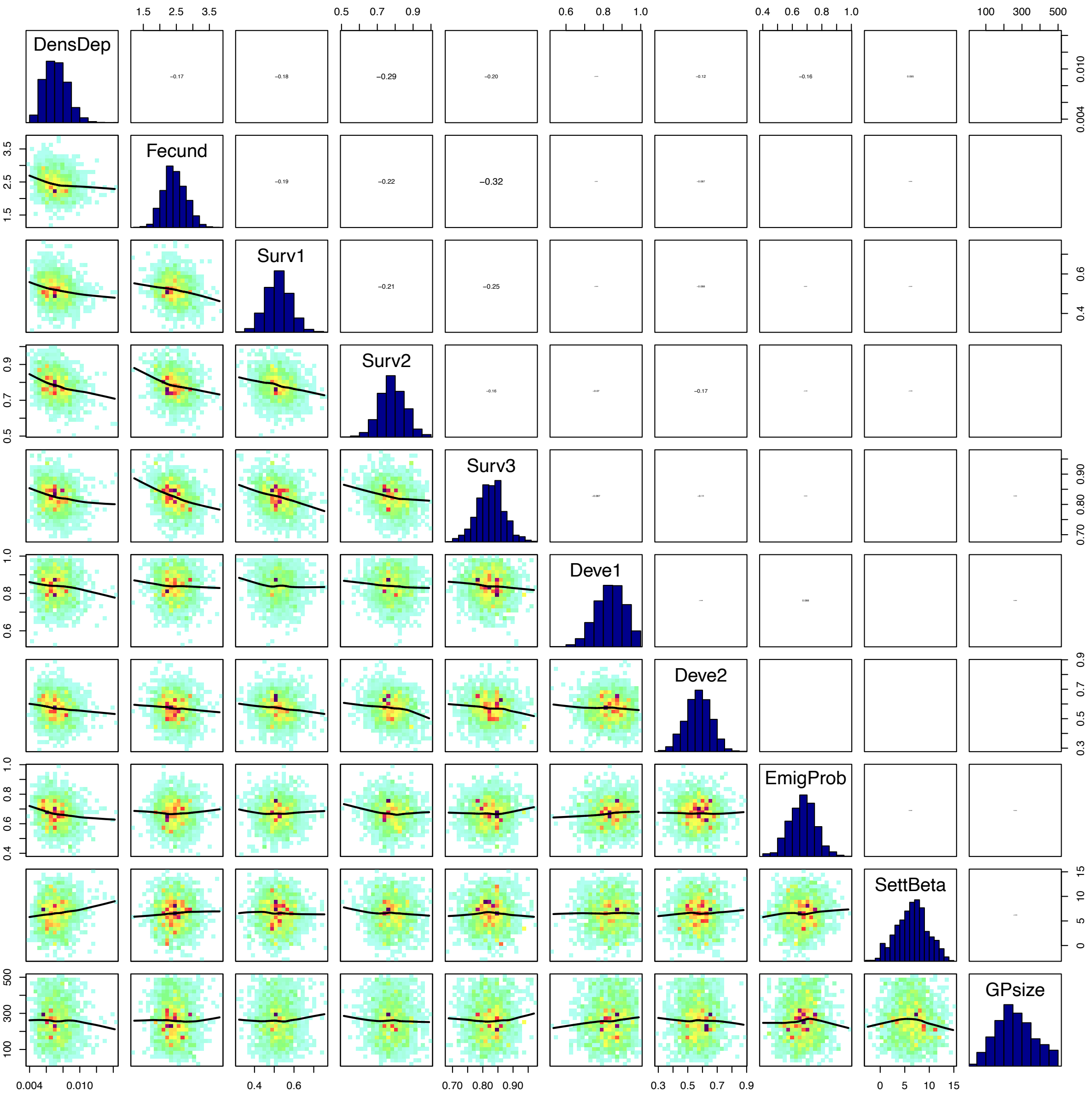


Figure S14: Pairwise posterior correlations

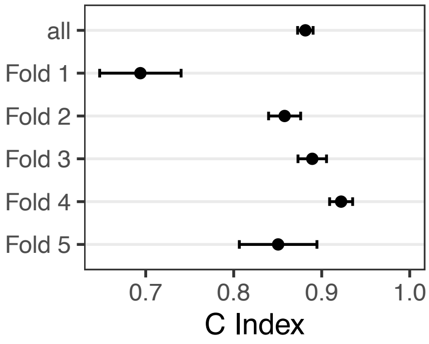


| Fold ID | C index | 95% CI |
|---|---|---|
| all | 0.88 | (0.87 - 0.89) |
| Fold 1 | 0.69 | (0.64 - 0.74) |
| Fold 2 | 0.86 | (0.84 - 0.88) |
| Fold 3 | 0.89 | (0.87 - 0.91) |
| Fold 4 | 0.92 | (0.91 - 0.93) |
| Fold 5 | 0.85 | (0.81 - 0.89) |

Figure S15: C-index of projections to the test data of the cross-validation. Given are the mean and 95%-confidence interval. ”all” refers to the whole MHB data set, where each observation was predicted from the particular calibration that had it in its test fold. The other rows provide the same info, but separate per fold (and calibration). Here, the c-index is calculated over all observation (site-year combination) independently.

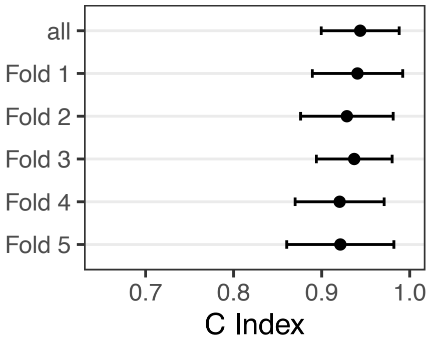


| Fold ID | C index | 95% CI |
|---|---|---|
| all | 0.94 | (0.90 - 0.98) |
| Fold 1 | 0.94 | (0.89 - 0.99) |
| Fold 2 | 0.93 | (0.88 - 0.98) |
| Fold 3 | 0.94 | (0.90 - 0.98) |
| Fold 4 | 0.92 | (0.87 - 0.97) |
| Fold 5 | 0.92 | (0.86 - 0.98) |

Figure S16: As Fig. S15, but the c-index is calculated over the time series of total abundance per fold.

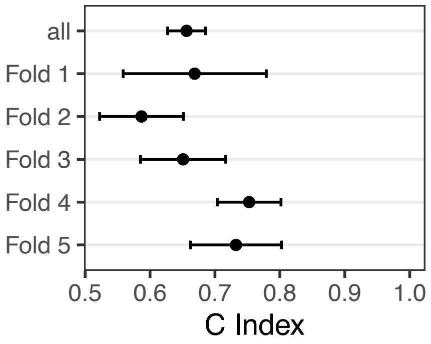


| Fold ID | C index | 95% CI |
|---|---|---|
| all | 0.66 | (0.63 - 0.69) |
| Fold 1 | 0.67 | (0.56 - 0.78) |
| Fold 2 | 0.59 | (0.53 - 0.65) |
| Fold 3 | 0.65 | (0.58 - 0.72) |
| Fold 4 | 0.75 | (0.70 - 0.80) |
| Fold 5 | 0.73 | (0.66 - 0.80) |

Figure S17: As Fig. S15, but the c-index is calculated for the 15% cells with highest variance in their local abundance time series.

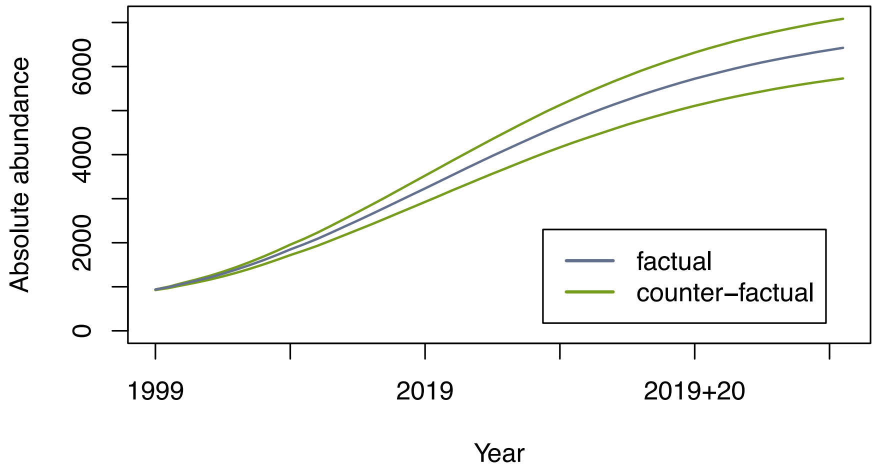


Figure S18: Sensitivity of the median posterior predictions to changes in the underlying habitat suitability maps. Shown are the time series of the median of total red kite abundance in Switzerland for the factual and two counter-factual scenarios. The factual scenario (blue line) is based on the observed environmental predictors and was used in the calibration. The two counter-factual scenarios (green lines) use habitat suitability maps in which the habitat suitability of each cell was increased or, respectively, decreased by five (out of a maximum of 100) points.

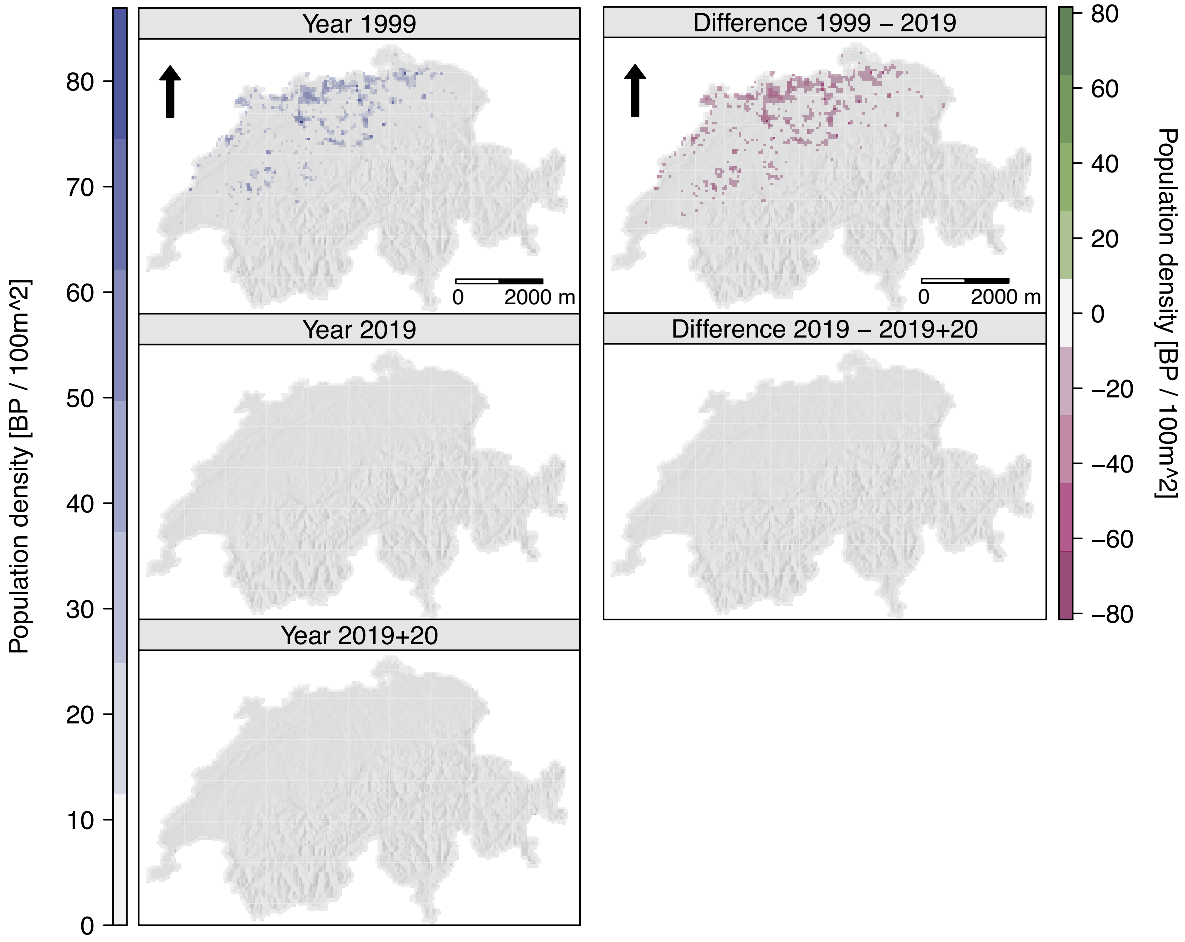


Figure S19: Mean prior predicted population densities for the years 1999 and 2019, and after 20 further years under constant conditions (left column) as well as the differences between those years (right column). The arrows indicate north.

### References

European Union (2022). *Copernicus Land Monitoring Service*. European Environment Agency (EEA).

Gelman, A. and Rubin, D. B. (1992). “Inference from Iterative Simulation Using Multiple Sequences”. *Statistical Science* 7.4, pp. 457–472. DOI: 10.1214/ss/1177011136.

Karger, D. N., Conrad, O., Böhner, J., Kawohl, T., Kreft, H., Soria-Auza, R. W., Zimmermann, N. E., Linder, H. P., and Kessler, M. (2017). “Climatologies at high resolution for the earth’s land surface areas”. *Scientific Data* 4, p. 170122. DOI: 10.1038/sdata.2017.122.

– (2018). *Data from: Climatologies at high resolution for the earth’s land surface areas*. DOI: 10.5061/dryad.kd1d4.

Katzenberger, J., Gottschalk, E., Balkenhol, N., and Waltert, M. (2019). “Long-term decline of juvenile survival in German Red Kites”. *Journal of Ornithology* 160.2, pp. 337–349. DOI: 10.1007/s10336-018-1619-z.

Knaus, P., Strebel, N., and Sattler, T. (2022). *The State of Birds in Switzerland 2022*. Sempach: Swiss Ornithological Institute.

Knaus, P., Jérôme, G., Sattler, T., Wechsler, S., Kéry, M., Strebel, N., and Antoniazza, S. (2018). *Schweizer Brutvogelatlas 2013-2016*. Sempach, Schweiz: Schweizerische Vogelwarte. ISBN: 978-3-85949-009-3.

Morris, M. D. (1991). “Factorial Sampling Plans for Preliminary Computational Experiments”. *Technometrics* 33.2, pp. 161–174. DOI: 10.1080/00401706.1991.10484804.

Nachtigall, W. (2008). “Der Rotmilan (Milvus milvus, L. 1758) in Sachsen und Südbrandenburg–Untersuchungen zu Verbreitung und Ökologie”. *Diss., Univ. Halle-Wittenberg*.

Nägeli, M., Scherler, P., Witczak, S., Catitti, B., Aebischer, A., Bergen, V. van, Kormann, U., and Grüebler, M. U. (2021). “Weather and food availability additively affect reproductive output in an expanding raptor population”. *Oecologia*. DOI: 10.1007/s00442-021-05076-6.

Newton, I., Davis, P., and Davis, J. (1989). “Age of first breeding, dispersal and survival of Red Kites Milvus milvus in Wales”. *Ibis* 131.1, pp. 16–21.

Schaub, M. (2012). “Spatial distribution of wind turbines is crucial for the survival of red kite populations”. *Biological Conservation* 155, pp. 111–118. DOI: 10.1016/j.biocon.2012.06.021.

Schmid, H., Luder, R., Naef-Daenzer, B., Graf, R., and Zbinden, N. (1998). *Schweizer Brutvogelatlas 1993–1996*. Sempach, Schweiz: Schweizerische Vogelwarte. ISBN: 978-3-9521064-5-7.

Schmid, H. and Volet, B. (2004). “Der Bestand des Rotmilans Milvus milvus im Winter 2002/03 in der Schweiz”. *Der Ornithologische Beobachter* 101, pp. 192–200.

Sergio, F., Tavecchia, G., Blas, J., Tanferna, A., and Hiraldo, F. (2021). “Demographic modeling to fine-tune conservation targets: importance of pre-adults for the decline of an endangered raptor”. *Ecological Applications* 31.3, e2266. DOI: https://doi.org/10.1002/eap.2266.

Vehtari, A., Gelman, A., Simpson, D., Carpenter, B., and Bürkner, P.-C. (2021). “Rank-Normalization, Folding, and Localization: An Improved $\hat{R}$ for Assessing Convergence of MCMC (with Discussion)”. *Bayesian Analysis* 16.2, pp. 667–718. DOI: 10.1214/20-BA1221.
