## Supplementary material for "Fitting individual-based models of spatial population dynamics to long-term monitoring data": ODD protocol for IBMs

#### S2 Appendix 2: ODD protocol of the red kite RangeShiftR model

This ODD protocol (Overview, Design concepts, and Details; Grimm el at. 2020) was generated by the RangeShiftR package. Its contents are part of the RangeShifter project user manual, written by Greta Bocedi, Stephen C. F. Palmer and Justin M. J. Travis from the University of Aberdeen, UK. It is available from the public github repository
`https://github.com/RangeShifter/RangeShifter-software-and-documentation/`.

#### Purpose and patterns

The RangeShiftR model is a single species, spatially-explicit and stochastic individual-based model (IBM), built around the integration of two fundamental components: population dynamics and dispersal behaviour. In this model of the red kite in Switzerland, population dynamics, represented as stage-structured, female-only population, with overlapping generations, and dispersal (explicitly modelled in its three phases of emigration, transfer and settlement, where the latter is accounting for context dependency as well as stage specificity) are played out on top of a gridded habitat suitability landscape.

### Entities, state variables and scales

#### Individuals

Individuals are the basic entities of the RangeShiftR model. Each individual has a unique ID number and here is defined by the following state variables:

- status
- initial (natal) and current location
- age and stage

#### Populations

Populations are defined by the individuals occupying a single cell and they represent the scale at which individuals interact and density-dependencies act. Populations are characterised by their size and location and the number in each stage class.

#### Landscape units

The model runs over a grid-based map.
Here, each cell stores a habitat suitability index, ranging from 0 to 100. This index is derived from a correlative species distribution model. Its value mediates the local strength of demographic density-dependence. Each cell is defined as suitable or not suitable for the red kite if its habitat suitability is above or equal to zero, respectively.

#### Spatial and temporal scales

The cell size (resolution) is 2000 m by 2000 m. The cell resolution represents the spatial scale at which the two fundamental processes of population dynamics and dispersal occur. This means that the density-dependency in the model (which here on reproduction and settlement) acts at the cell scale and cells are the single step unit for the discrete movement model used in the transfer phase. There are three distinct temporal scales: The highest-level represents years and encompass a full model cycle. The intermediate scale is the species' reproductive season. The model simulates one reproductive season per year. Finally, the smallest time scale is represented by the number of steps that dispersers take during the movement phase of dispersal. This is determined by the maximum number of steps.

### Process overview and scheduling

At the beginning of each year, reproduction is the first process to be modelled. Reproduction is followed by natal dispersal. After each reproductive season, survival and successive development of all the stages are modelled. Aging occurs at the end of the year.

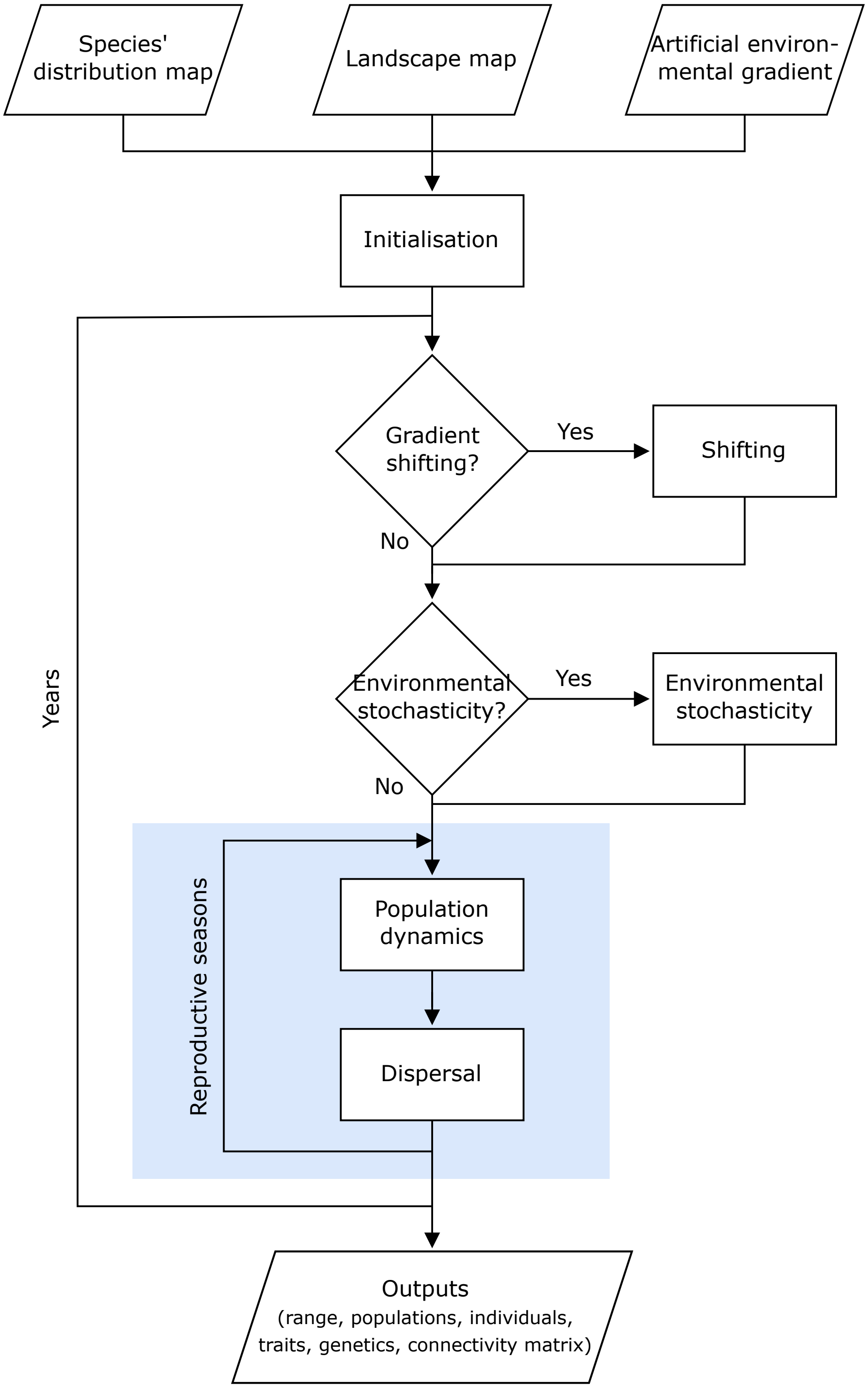


Figure S1: General model workflow and schedule. The model core (highlighted in blue) is expanded in Fig. S2.

### Design concepts

#### Basic principles

##### Population dynamics

Demographic stochasticity is fundamentally important for the dynamics of populations that are naturally small or have declined to low abundances owing to anthropogenic pressures. Additionally, inter-individual variability within populations can have a major influence on dynamics. Modelling stochastic events that happen to individuals is crucial for avoiding systematic overestimation of population viability or rate of spread (Clark et al. 2001, Kendall and Fox 2003, Robert et al. 2003, Grimm and Railsback 2005, Jongejans et al. 2008, Travis et al. 2011). Thus, population dynamics in RangeShiftR were constructed to be fully individual-based and stochastic. Each reproductive individual produces a discrete number of offspring sampled from a Poisson distribution with a mean that is influenced by the species' demographic parameters and the local population density.

##### Definition of populations

The cell is the scale at which processes such as population dynamics and dispersal act. The individuals present in a cell define a distinct population, and density-dependencies for reproduction and settlement both operate at this scale. Even in the case where two habitat cells are adjacent, they still hold separate populations.

##### Population structure

Implementing overlapping generations which are stage-structured is the appropriate choice for species in which generations can overlap and individuals can be classified in different stages (e.g. immature vs. breeding individuals) differing in their demographic parameters. Individuals are characterized by their age and stage. Each stage has a certain fecundity, survival and probability of developing to the next stage. The parameters are provided through classical transition matrices (Caswell 2001). However, in RangeShiftR, these are not multiplied with the population vector as is typical for matrix models but, instead, the parameters are applied stochastically in an individual-based manner. Presenting the demographic parameters in the standard matrix notation, is meant to ease parameterisation Bocedi et al. (2014), as most population modellers are used to matrix models. It has the further important benefit of helping bridging the gap between analytical models and IBMs, the joint use of which has considerable potential, especially for improving modelling for conservation (Travis et al. 2011).

##### Dispersal

Dispersal is defined as movement leading to spatial gene flow, and it typically involves three phases: emigration, transfer and settlement (Stenseth and Lidicker 1992, Clobert et al. 2001, Bowler and

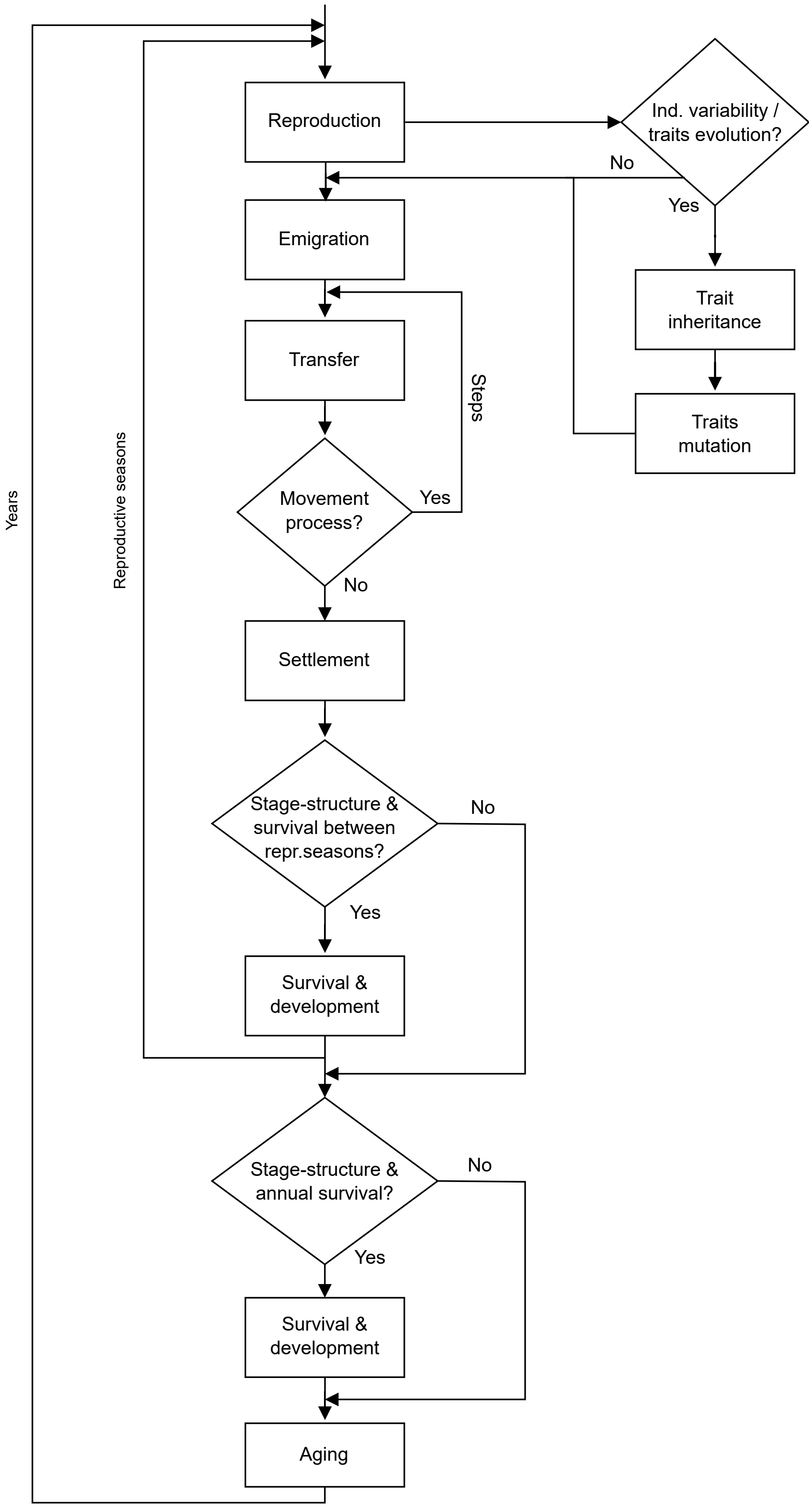


Figure S2: General flowchart of the core model.

Benton 2005, Ronce 2007). The key role of dispersal in species persistence and responses to environmental change is increasingly recognized (Travis et al. 2014). Moreover, the importance of modelling dispersal as a complex process, explicitly considering its three phases, each of which has its own mechanisms and costs, has been recently highlighted (Travis et al. 2012, 2014, Bonte et al. 2012). The implementation of the dispersal process in RangeShiftR is based on these recent frameworks and the substantial dispersal theory that has been developed so far (Clobert et al. 2012).

**Emigration** Emigration is the first phase of dispersal. Emigration itself can be a complex process determined by multiple proximate and ultimate causes. Multiple emigration strategies can be present across the species' range, inside a single population or even within the same individual in form of plastic emigration behaviour.

The theory on emigration accounts for context dependencies, plasticity and inter-individual variability in emigration strategies. Much work has been conducted to understand the role of density dependence in emigration (Travis et al. 1999, Metz and Gyllenberg 2001, Poethke and Hovestadt 2002, Matthysen 2005, Kun and Scheuring 2006, Chaput-Bardy et al. 2010, De Meester and Bonte 2010).

In RangeShiftR, emigration is modelled as the probability that an individual will leave its natal patch during the present year. See the description of the submodel for further details.

**Transfer** Transfer is the second phase of dispersal, and consists of the movement of an individual starting from when it emigrates from its natal patch and ending with settlement in another patch or with dispersal mortality. The main components of this phase are the individual movement ability and navigation capacity in response to the characteristics of the environment. The interaction between these components and their associated costs will determine the distance moved, the movement path and the chance of surviving the transfer phase.

Understanding and modelling how species move is not a simple task, and, perhaps more than for the other phases of dispersal, much effort has been spent in two separate and not always interacting fields: dispersal ecology (Travis et al. 2010) and movement ecology (Nathan et al. 2008). While the former seeks to understand movements as a part of the dispersal process and has often described transfer with phenomenological dispersal kernels (but see recent developments in fitting mechanistic kernels: (Schurr 2012)), the latter is more focused on understanding the mechanisms of the movement process itself, even though recent emphasis has been put on the consequences of movements for population dynamics (Morales et al. 2010). Modelling dispersal in IBMs needs to draw from both fields.

It is increasingly acknowledged that individual movements within and between habitat patches, and consequently also population dynamics, are strongly affected by the behavioural and physical traits of individuals and by the landscape structure and composition (Morales and Ellner 2002, Hawkes 2009, Stevens and Coulon 2012, Baguette et al. 2013). This has led to the development of mechanistic models where movement behaviour and its interaction with the environment is explicitly described (Nathan et al. 2008, Revilla and Wiegand 2008, Morales et al. 2010, Palmer et al. 2011, Pe'er et al. 2011). The classical method to represent individuals' movements mechanistically is to use a random walk (Codling et al. 2008), or its diffusion approximation, assuming that individuals are moving randomly in a homogeneous landscape and that they are all following the same rules. From this basis, there have been recent developments in diffusion models for including landscape heterogeneity and some behavioural responses, like reaction to habitat boundaries, directly derived from empirical data through state-space models (Ovaskainen and Cornell 2003, Ovaskainen 2004, Patterson et al. 2008, Ovaskainen et al. 2008, Ovaskainen and Crone 2009, Zheng et al. 2009). Yet, these models do not account for individual variability or for many behavioural components including memory, perceptual range and movement modes. Despite this simplicity, diffusion models, and especially their recent developments, can still be satisfactory at large temporal and spatial scales and serve as a null hypothesis against which to test more complex movement models. Moreover they can provide basis for building blocks for population dynamics models.

Mechanistic IBMs allow extending the "random paradigm" by incorporating behavioural elements that are likely to be crucial in affecting species' spatial dynamics (Lima and Zollner 1996, Baguette and Van Dyck 2007, Knowlton and Graham 2010, Shreeve and Dennis 2011). These elements can be assigned into six main categories: (i) the switching between different movement modes [for example foraging within the home range vs. dispersal (Fryxell et al. 2008, Delattre et al. 2010, Pe'er et al. 2011)]; (ii) the individuals' perceptual range (Zollner and Lima 1997, Olden et al. 2004, Gardner and Gustafson 2004, Zollner and Lima 2005, Vuilleumier et al. 2006, Vuilleumier and Metzger 2006, Pe'er and Kramer-Schadt 2008, Palmer et al. 2011); (iii) the use of information in movement choices (Clobert et al. 2009) and the memory of previous experience (Smouse et al. 2010); (iv) the influence of habitat fragmentation and matrix heterogeneity on movement behaviours (Ricketts 2001, Vandermeer and Carvajal 2001, Schtickzelle and Baguette 2003, Revilla et al. 2004, Wiegand et al. 2005, Fahrig 2007, Dover and Settele 2009); (v) the individual responses to habitat boundaries (Schultz and Crone 2001, Morales 2002, Merckx et al. 2003, Ovaskainen 2004, Stevens et al. 2006, Pe'er et al. 2011); and (vi) the period of activity (Revilla et al. 2004) and the time scale of movements (Lambin et al. 2012).

A general framework for a mechanistic representation of movements has been outlined by Nathan et al. (2008), who identified four basic components: the internal state of the individual (why does it move?), its motion capacities (how does it move?), its navigation capacities (when and where does it move?) and external factors that affect the movement. This framework allows us, starting from individual movements, and taking into account individual variability, to predict movement patterns over large temporal and spatial scales and potentially to scale up to populations, communities, ecosystems and to multi-generation / evolutionary processes (Holyoak et al. 2008). The ultimate limitation is likely to be the quantity and the type of data needed to parameterize this /these kind of models; therefore, the challenge is to understand which level of detail is needed to make reliable projections in different contexts and for different purposes (Lima and Zollner 1996, Morales et al. 2010).

Movement behaviours during the transfer phase are a core component of the dispersal strategy of an individual, and therefore they come under selection and they can evolve (Merckx et al. 2003, Fahrig 2007, Hawkes 2009, Travis et al. 2012). Ultimately, it is the evolution of movement behaviours that leads to what we consider the evolution of dispersal kernels. A handful of theoretical studies have so far explored the evolution of movement rules. For example, it has been shown how the landscape composition and configuration, in interaction with the ecology of the species, can affect the evolution of movement patterns, such that the greater the costs of dispersal the more highly correlated are the emerging walks (Heinz and Strand 2006, Bartoń et al. 2009). Moreover, straighter movement paths (Phillips et al. 2010) and riskier strategies seem to be selected during range expansion in such a way that the rate of expansion is maximized at the expense of the survival probability of the single individual (Bartoń et al. 2012).

Here, transfer is implemented as a strongly correlated random walk. This submodel is fully individual-based and explicitly describes the movement behaviour of individuals with a level of detail, and hence parameters, which is probably close to the most parsimonious for a mechanistic movement model. However, it facilitates considerably increasing the complexity and realism with which the transfer phase is modelled.

See the submodel description for further details.

**Settlement** Settlement, or immigration, is the last phase of dispersal, when the organism stops in a new cell or patch of breeding habitat. This phase is determined by a suite of strategies, behaviours and reaction norms that lead individuals to the decision to stop in a particular place. Habitat selection, mate finding and density dependence are probably three of the main processes involved, but not the only ones. Like emigration, settlement is a complex process affected by multiple criteria including inter-individual variability and context dependencies. It can be influenced by the causes and mechanisms of the previous phases of dispersal (Clobert et al. 2009) and it has associated specific costs (Bonte et al. 2012), which can also feed back to the previous phases (Le Galliard et al. 2012).

As for the previous phases, the use of different sources of abiotic and biotic information is likely to be crucial in the settlement decision, for which evidence is now accumulating. For example, studies have demonstrated that in some species, dispersing individuals exhibit a preference for habitat that is similar to the natal one, philopatry being a stronger predictor of habitat preferences for settlement than intrinsic habitat quality (Haughland and Larsen 2004, Stamps and Blozis 2006, Stamps et al. 2009). Conspecific density and performance have also been demonstrated to be important cues for settlement decisions (conspecific attraction), because they can provide a rapid approximation of the habitat quality (Stamps 1988, Doligez et al. 2004, Fletcher 2007, Vercken et al. 2012, Clotuche et al. 2013).

From the theoretical point of view, much work has been conducted on habitat selection during settlement decisions and its consequences for species' population dynamics and spatial genetic structure. The basic assumption is that individuals are expected to select habitat patches where their expected fitness is greater than the one expected in the natal patch, weighted by the costs of searching (Ruxton and Rohani 1998, Stamps 2001, Baker and Rao 2004, Stamps et al. 2005, Armsworth and Roughgarden 2008, Bonte et al. 2012). Recently the idea of 'matching habitat choice' has been proposed, for which individuals aim to settle where the environment best matches with their phenotype. This process, expected to be more important for species with limited phenotypic plasticity, can have important implications for processes such as local adaptation, adaptive peak shifts and evolution of niche width, and speciation (Edelaar et al. 2008). Other factors affecting settlement such as density dependence (Poethke et al. 2011), conspecific attraction (Fletcher 2006) or mate finding (Gilroy and Lockwood 2012), their evolution and their consequences on species' responses to environmental changes, have been much less theoretically investigated.

This model incorporates density-dependent settlement rules. Dispersing individuals are not allowed to settle in their natal cell or a non-suitable cell and settlement probability is dependent on the size of the target cell's local population. For more details, see the description of the submodel.

##### Dispersal mortality

Dispersal is often a costly process for an organism (Bonte et al. 2012) and, in some cases, a dispersing individual may suffer mortality. Obtaining a sensible representation of dispersal requires that these mortality costs are described appropriately and, for this, it is important to recognize how dispersal mortality is incorporated in RangeShiftR. First, dispersal mortality can arise as a result of individuals failing to reach suitable habitat. For example, some individuals may fail to find suitable habitat before they use up a maximum number of movement steps. In this first case, dispersal mortality clearly depends on the proportion of suitable habitat in the landscape and will increase as the availability of habitat declines.

A second source of dispersal mortality can be a per-step probability of mortality. This can be useful for representing mortality risks that increase with distance or time spent travelling.

For more details, see the description of the submodel.

#### Emergence

Species range distributions emerge from the population dynamics and the dispersal behaviour of the individuals.

#### Adaptation

Not applicable.

#### Objectives

Not applicable.

#### Learning

Not applicable.

#### Prediction

Not applicable.

#### Sensing

Dispersing individuals can perceive the density in their current cell and potentially decide to settle if local density is low or to continue dispersing if it is high.

#### Interaction

No interaction is considered in the model.

#### Stochasticity

Demographic stochasticity is explicitly included in all processes.

#### Collectives

Not applicable

#### Observation

##### Populations

The population output consists of a raster stack with layers for each year and for each simulation replicate. The layers are gridded maps of the simulation domain in which each cell holds the number of breeding adults in its population.

### Initialization

#### Initialization of landscape

Landscapes are initialized according to input raster maps, that are generated as projections from a correlative species distribution model. Thus, the habitat suitability in each cell is assumed as the modelled red kite occurrence probability in this cell. The landscape is re-initialised each year as the species distribution model is evaluated with the current predictors in each year and thus the habitat suitabilities slightly change. The demographic density-dependence of each cell is given as the maximum density-dependence (denoted $1/b$) times the cell's habitat suitability index in percent.

#### Initialization of populations

Populations are initialized from a data frame that lists the initial individuals with their given age, stage, and natal (i.e. initial) cell. This initial condition is generated from a Poisson model that takes into account the local habitat suitability and spatial correlation. The distributions of stages and ages of initialised individuals are calculated from the transition matrix as the quasi-equilibrium distributions.

### Input data

The model requires the following input data:

- Landscape maps as 'raster' objects in R.
  Several maps may be given if the landscape shall change at specific years during the simulation. The grids need to contain habitat suitability values for each cell. Each cell in the landscapes is assigned a continuous percentage value between 0.0 and 100.0 (of the maximum density-dependence $1/b$). There are no explicit habitat or land-cover types. This allows integrating different methods for calculating the habitat suitability for a given species. For example, qualities can result from different methods of suitability modelling, which incorporate multiple variables like habitat types, elevation, climate, etc. A linear relationship between the habitat quality and the actual density-dependence of the population dynamics is assumed. Therefore, the quality should be scaled accordingly in case of a curvilinear relationship. Any part of the original landscape which was a 'no-data' region (e.g. the sea or land beyond a study area boundary) must remain in that state for the whole simulation.
- A list of initial individuals.
  The population is initialised according to a list of initial individuals (of given age and stage) in specified cells and years. This option allows simulation of a reintroduction scenario.

### Submodels

#### Landscape

Dynamic landscapes are imported as a set of gridded maps that come into effect at specified years during the simulation. Each gridded map holds continuous values in its cells, ranging from 0.0 to 100.0, that represent percentages of habitat cover or quality. Thus, at a maps change some populations may be extirpated (all individuals either die or have an immediate opportunity to disperse) if suitable habitat is lost, and new populations may arise from colonisation of newly suitable areas.

In addition, the demographic density-dependence $1/b$ is given in units of the number of individuals per hectare. The given density dependence is interpreted as the maximum stregnth of demographic density-dependence reached in cells with 100% habitat. All other cells hold the respective fraction of the stregnth of density-dependence.

#### Reproduction

At the beginning of each year, reproduction is the first process to be modelled. The model simulates one reproductive season per year. Reproduction is followed by dispersal. After each reproductive season, survival and successive development of all the stages are modelled. Aging occurs at the end of the year.

Generations overlap and individuals are classified in different stages (e.g. immature vs. breeding individuals) differing in their demographic parameters. Individuals are characterized by their age and stage. Each stage has a certain fecundity $\phi$, survival $\sigma$ and probability of developing to the next stage $\gamma$. The parameters are provided through classical transition matrices (Caswell 2001). However, these are not multiplied with a population state vector as is typical for matrix models but, instead, the parameters are applied stochastically in an individual-based manner. At each reproductive season, two parameters control the likelihood that each individual / female reproduces:

- First, it is determined whether a reproductively mature female is a potential reproducer. Only those mature females that are able to reproduce, are potential breeders.
- Potential breeders all reproduce with a set probability $\phi$.

In this model of the red kite, a female-only model was implemented. It models the three stages of juveniles, sub-adults and breeding adults. To allow offspring to developing to the next stage in the same year without losing an explicit offspring stage in which post-natal dispersal can happen, an explicit newborn stage (stage 0) is added, for example in a generic 3-stage model:

$$A = \begin{pmatrix} 0 & \phi_1 & \phi_2 & \phi_3 \\ 1 & \sigma_1 & 0 & 0 \\ 0 & \gamma_{1-2} & \sigma_2 & 0 \\ 0 & 0 & \gamma_{2-3} & \sigma_3 \end{pmatrix}$$

Newborn have to develop to stage 1 in the same year they are born. It is important to note that juvenile mortality can be accounted for in two ways. Either it is included in adult fecundity $\phi$ (by appropriately reducing its value), and $\gamma_{0-1}$ is equal to 1.0. This is how it is typically accounted for in matrix models. Or, alternatively, $\phi$ is equal to the true maximum fecundity and $\gamma_{0-1}$ is less than 1.0. In any case, $\sigma_0$ needs to be zero. Only the first approach allows straightforward direct comparison with standard analytical matrix models. The parameters in the matrix are used in a stochastic way at the individual level. Each individual/female at stage 3, if it reproduces, produces a number of offspring given by Poisson($\phi_3$).

Reproduction is density- and stage-dependent. Following Neubert & Caswell (2000), density and stage dependence in fecundity is implemented as an exponential decay:

$$\phi_i = \phi_{0,i} * e^{-b \sum_{j=1}^{S} \omega_{ij} N_{j,t}}$$

where $\phi_i$ is the fecundity of stage $i$, $\phi_{0,i}$ is its maximum fecundity at low densities, $b$ is the strength of density dependence, $S$ indicates the number of stages and $\omega_{ij}$ is contribution of stage $j$ to the density dependence in the fecundity of stage $i$ and $N_{j,t}$ the number of individuals in stage $j$ at time $t$ (e.g. (Caswell et al. 2004)).

#### Survival & Development

Reproduction is first followed by dispersal. Only then survival and successive development of all the stages are modelled.

Here, survival is neither density- nor stage- dependent. Bernoulli trials $Bern(\sigma)$ determine if an individual survives or not.

Likewise, development is neither density- nor stage- dependent. Bernoulli trials $Bern(\gamma)$ determine if an individual develops to the next stage or not.

#### Emigration

Emigration is the first phase of dispersal. It is modeled as the probability $e$ that an individual will leave its natal patch during the present year. If a stage having non-zero $e$ can last for more than one year, an individual has multiple opportunities to emigrate, each with probability $e$, and hence the realised overall emigration rate will be larger than $e$. Here, the emigration probability $e$ is density-independent but stage-specific, so that only juveniles and sub-adults emigrate.

#### Transfer

Transfer is the second phase of dispersal, and consists of the movement of an individual starting from when it emigrates from its natal cell and ending with settlement in another cell or mortality. The main components of this phase are the individual movement ability and navigation capacity in response to the characteristics of the environment. The interaction between these components and their associated costs will determine the distance moved, the movement path and the chance of surviving the transfer phase.

The movement model is fully individual-based and explicitly decribes the movement behaviour of individuals with a level of detail, and hence parameters, which is probably close to the most parsimonious for a mechanisctic movement model.

For this model, a simple correlated random walk is used. This transfer submodel is implemented in continuous space on the top of the landscape grid. Individuals take steps of a constant step length, the direction is sampled from a wrapped Cauchy distribution having a correlation parameter $\rho$ in the range 0 to 1 (Zollner and Lima 1999, Bartoń et al. 2009). All individuals take each step simultaneously.

#### Settlement

Settlement, or immigration, is the last phase of dispersal, when the organism stops in a new cell of breeding habitat. Dispersing individuals are not allowed to settle in their natal cell.

At each step, made simultaneously by all dispersing individuals, each evaluates their current cell for the possibility of settling. The individual decides to stop if there is suitable habitat; this is a necessary condition. Additionally, the settlement decision is density-dependent. The individual has a probability $p_s$ of settling in the cell $i$, given by:

$$p_s = \frac{S_0}{1 + e^{-(bN_i - \beta_i) * \alpha_s}}$$

Here, $N_i$ is the number of individuals of the cell $i$, $b$ represents the strength of density dependence used for the population dynamics, $S_0$ is the maximum settlement probability, $\beta_s$ is the inflection point and $\alpha_s$ is the slope of the function.

To avoid having individuals moving perpetually because they cannot find suitable conditions to settle, the model includes a maximum number of steps. The maximum number of steps defines the maximum time length of the transfer period. When an individual reaches the maximum number of steps, it stops where it is regardless of the suitability of the location. In the next season, if still alive, it will continue to move again.

#### Aging

Aging occurs at the end of each year.

### Model parameter settings

Table S1: Simulation parameters

| Parameter | Description | Values |
|---|---|---|
| Year | Number of simulated years | 24 |
| Replicates | Number of simulation iterations | 20 |
| Absorbing | Whether non-valid cells lead to direct mortality of the individual during transfer | FALSE |
| LocalExt | Local extinction | FALSE |
| EnvStoch | Environmental stochasticity (0: none, 1: global, 2: local) stochasticity acts (0: growth rate/fecundity, 1: demographic density dependence) | 0 |
| OutIntRange | Output of range file | 0 |
| OutIntOcc | Output of occupancy file | 0 |

| Parameter | Description | Values |
|---|---|---|
| OutIntPop | Output of population file | 1 |
| OutIntInd | Output of individual file | 0 |
| OutIntConn | Output of connectivity file | 0 |
| OutIntPaths | Output of SMS paths file | 0 |
| OutStartPop | Starting year for output population file | 2 |
| OutStartInd | Starting year for output individual file | 0 |
| OutStartConn | Starting year for output connectivity file | 0 |
| OutStartPaths | Starting year for output SMS paths file | 0 |
| SMSHeatMap | Output SMS heat map raster file | FALSE |

Table S2: Landscape parameters

| Parameter | Description | Values |
|---|---|---|
| Landscape file | Landscape map(s) | habitatmaps |
| Resolution | Resolution in meters | 2000 |
| HabPercent | Whether habitat types/codes or habitat cover/quality | TRUE |
| NHabitats | Number of different habitat codes | 2 |
| K_or_DensDep | Demographic density-dependence | 0.006 |
| PatchFile | Filename(s) of the patch map(s) | NULL |
| CostsFile | Filename(s) of the SMS cost map(s) | NULL |
| Dynami-cLandYears | Years of landscape changes | 0, 1, 2, 3, 4, 5, 6, 7, 8, 9, 10, 11, 12, 13, 14, 15, 16, 17, 18, 19 |
| InitIndsFile | Filename of the species initial distribution list | NULL |
| InitIndsList | Species initial distribution data frame | createInitIndsFile_Pois() |

Table S3: Demography parameters

| Parameter | Description | Values |
|---|---|---|
| Stages | Number of life stages | 4 |
| TransMatrix | Transition matrix | $\begin{bmatrix} 0 & 0 & 0 & 1.65 \\ 1 & 0.084 & 0 & 0 \\ 0 & 0.336 & 0.306 & 0 \\ 0 & 0 & 0.374 & 0.8 \end{bmatrix}$ |
| | Defines the development probabilities from each stage into the next as well as the respective survival probabilities and fecundities | |
| MaxAge | Maximum age in years | 12 |
| MinAge | Ages which an individual in stage i-1 must already have reached before it can develop into the next stage i. | 0, 0, 0, 0 |
| RepSeasons | Number of potential reproduction events per year | 1 |
| RepInterval | Number of reproductive seasons which must be missed following a reproduction attempt before another reproduction attempt may occur | 0 |
| PRep | Probability of reproducing in subsequent reproductive seasons | 1 |
| SurvSched | Scheduling of survival (0: at reproduction, 1: between reproductive events, 2: annually) | 1 |
| FecDensDep | whether density dependent fecundity probability is modelled | TRUE |
| DevDensDep | Whether density dependent development probability is modelled | FALSE |
| SurvDensDep | Whether density dependent | FALSE |

| Parameter | Description | Values |
|---|---|---|
| | survival probability is modelled | |
| DevDensCoeff | Relative density dependence coefficient for development | 1 |
| SurvDensCoeff | Relative density dependence coefficient for survival | 1 |
| FecStageWtsMatrix | Stage-dependent weights in density dependence of fecundity | $\begin{bmatrix} 0 & 0 & 0 & 0 \\ 0 & 0 & 0 & 0 \\ 0 & 0 & 0 & 0 \\ 0 & 0 & 1 & 1 \end{bmatrix}$ |
| DevStageWtsMatrix | Stage-dependent weights in density dependence of development | Not selected. |
| SurvStageWtsMatrix | Stage dependent weights in density dependence of survival | Not selected. |
| PostDestrictn | Whether individuals of a population die (FALSE) or disperse (TRUE) if its patch gets destroyed | FALSE |
| ReproductionType | Decribes the reproduction type (0: asexual/only female; 1: simple sexual model; 2: sexual model with explicit mating system) | 0 |

Table S4: Dispersal parameters

| Process | Parameter | Description | Values |
|---|---|---|---|
| Emigration | EmigProb | Emigration probabilities per stage | $\begin{bmatrix} 0 & 0 \\ 1 & 0.64 \\ 2 & 0 \\ 3 & 0 \end{bmatrix}$ |
| | SexDep | Sex-dependent emigration probability? | FALSE |

| Process | Parameter | Description | Values |
|---|---|---|---|
| | StageDep | Stage-dependent emigration probability? | TRUE |
| | DensDep | Density-dependent emigration probability? | FALSE |
| | UseFullKern | Shall the emigration probability be derived from dispersal kernel? | FALSE |
| Transfer | Model | Type of transfer model | Correlated Random Walk |
| | StepLength | Step length given in meters | 2000 |
| | Rho | Correlation parameter | 0.85 |
| | StraightenPath | Straighten path after decision not to settle in a patch? | FALSE |
| | StepMort | Per-step mortality probability | 0.00 |
| Settlement | DensDep | Density-dependent settlement requirements? | TRUE |
| | SexDep | Sex-dependent settlement requirements? | FALSE |
| | StageDep | Stage-dependent settlement requirements? | FALSE |
| | Settle | Settlement probability parameters for all stages/sexes. | $\begin{bmatrix} 0.85 & -1 & 1.67 \end{bmatrix}$ |
| | MinSteps | Minimum number of steps | 0 |
| | MaxSteps | Maximum number of steps | 10 |
| | MaxStepsYear | Maximum number of steps per year | 5 |
| | FindMate | Mating requirements to settle? | FALSE |

Table S5: Initialisation parameters

| Parameter | Description | Values |
|---|---|---|
| InitType | Type of initialisation<br>(0: free initialisation according to habitat map,<br>1: from loaded species distribution map,<br>2: from initial individuals list file) | 2 |

| Parameter | Description | Values |
|---|---|---|
| | (0: random in given number of cells, | |
| | 1: all suitable cells/patches) | |
| | species distribution map (0: all suitable cells within | |
| | all distribution presence cells, | |
| | 1: all suitable cells within given | |
| | number of randomly chosen presence cells) | |
| InitDens | Number of individuals seeded in each cell/patch | 0 |
| | (0: at demographic density dependence, | |
| | 1: at half of the demographic density dependence, | |
| | 2: according to quasi-equilibrium distribution) | |
| | (0: minimum age for the respective stage, | |
| | 1: random age between the minimum | |
| | and maximum age for the respective stage, | |
| | 2: according to a quasi-equilibrium distribution) | |
| InitFreezeYear | Year until which species is | 0 |
| | confined to its initial range limits | |
| RestrictRows | Number of rows at northern | 0 |
| | front to restrict range. | |
| RestrictFreq | Frequency in years at which | 0 |
| | range is restricted to northern front. | |
| FinalFreezeYear | The year after which species is | 0 |
| | confined to its new, current range limits, | |
| | after a period of range expansion. | |

#### References


Armsworth, Paul R. and Roughgarden, Joan E. 2008. The Structure of Clines with Fitness-Dependent Dispersal. - The American Naturalist 172: 648–657.

Baguette, M. and Van Dyck, H. 2007. Landscape connectivity and animal behavior: functional grain as a key determinant for dispersal. - Landscape Ecol 22: 1117–1129.

Baguette, M. et al. 2013. Individual dispersal, landscape connectivity and ecological networks. - Biological Reviews 88: 310–326.

Baker, M. B. and Rao, S. 2004. Incremental Costs and Benefits Shape Natal Dispersal: Theory and Example with Hemilepistus reaumuri. - Ecology 85: 1039–1051.

Bartoń, K. A. et al. 2009. The evolution of an “intelligent” dispersal strategy: biased, correlated random walks in patchy landscapes. - Oikos 118: 309–319.

Bartoń, K. A. et al. 2012. Risky movement increases the rate of range expansion. - Proceedings of the Royal Society B: Biological Sciences 279: 1194–1202.

Bocedi, G. et al. 2014. RangeShiftR: a platform for modelling spatial eco-evolutionary dynamics and species’ responses to environmental changes. - Methods in Ecology and Evolution 5: 388–396.

Bonte, D. et al. 2012. Costs of dispersal. - Biol Rev Camb Philos Soc 87: 290–312.

Bowler, D. E. and Benton, T. G. 2005. Causes and consequences of animal dispersal strategies: relating individual behaviour to spatial dynamics. - Biol Rev Camb Philos Soc 80: 205–225.

Caswell, H. 2001. Matrix Population Models: Construction, Analysis, and Interpretation. - Oxford University Press.

Caswell, H. et al. 2004. Sensitivity analysis of equilibrium in density-dependent matrix population models. - Ecology Letters 7: 380–387.

Chaput-Bardy, A. et al. 2010. Condition and Phenotype-Dependent Dispersal in a Damselfly, Calopteryx splendens. - PLOS ONE 5: e10694.

Clark, J. S. et al. 2001. Invasion by extremes: population spread with variation in dispersal and reproduction. - Am Nat 157: 537–554.

Clobert, J. et al. 2001. Dispersal. - Oxford University Press.

Clobert, J. et al. 2012. Dispersal Ecology and Evolution - Oxford University Press.

Clobert, J. et al. 2009. Informed dispersal, heterogeneity in animal dispersal syndromes and the dynamics of spatially structured populations. - Ecol Lett 12: 197–209.

Clotuche, G. et al. 2013. Settlement decisions by the two-spotted spider mite Tetranychus urticae. - Comptes Rendus Biologies 336: 93–101.

Codling, E. A. et al. 2008. Random walk models in biology. - J R Soc Interface 5: 813–834.

De Meester, N. and Bonte, D. 2010. Information use and density-dependent emigration in an agrobiont spider. - Behavioral Ecology 21: 992–998.

Delattre, T. et al. 2010. Dispersal mood revealed by shifts from routine to direct flights in the meadow brown butterfly Maniola jurtina. - Oikos 119: 1900–1908.

Doligez, B. et al. 2004. Availability and Use of Public Information and Conspecific Density for Settlement Decisions in the Collared Flycatcher. - Journal of Animal Ecology 73: 75–87.

Dover, J. and Settele, J. 2009. The influences of landscape structure on butterfly distribution and movement: a review. - J Insect Conserv 13: 3–27.

Edelaar, P. et al. 2008. Matching Habitat Choice Causes Directed Gene Flow: A Neglected Dimension in Evolution and Ecology. - Evolution 62: 2462–2472.

Fahrig, L. 2007. Non-optimal animal movement in human-altered landscapes. - Functional Ecology 21: 1003–1015.

Fletcher, R. J. 2007. Species interactions and population density mediate the use of social cues for habitat selection. - J Anim Ecol 76: 598–606.

Fryxell, J. M. et al. 2008. Multiple movement modes by large herbivores at multiple spatiotemporal scales. - PNAS 105: 19114–19119.

Gardner, R. H. and Gustafson, E. J. 2004. Simulating dispersal of reintroduced species within heterogeneous landscapes. - Ecological Modelling 171: 339–358.

Gilroy, J. J. and Lockwood, J. L. 2012. Mate-Finding as an Overlooked Critical Determinant of Dispersal Variation in Sexually-Reproducing Animals. - PLOS ONE 7: e38091.

Grimm, V. and Railsback, S. F. 2005. Individual-based Modeling and Ecology. - Princeton University Press.

Grimm, V., Railsback, S. F., Vincenot, C. E., Berger, U., Gallagher, C., Deangelis, D. L., Edmonds, B., Ge, J., Giske, J., Groeneveld, J., Johnston, A. S. A., Milles, A., Nabe-Nielsen, J., Polhill, J. G., Radchuk, V., Rohwäder, M. S., Stillman, R. A., Thiele, J. C., and Ayllón, D. 2020. The ODD protocol for describing agent-based and other simulation models: A second update to improve clarity, replication, and structural realism. - Journal of Artificial Societies and Social Simulation, 23(2), Article 2.

Haughland, D. L. and Larsen, K. W. 2004. Exploration correlates with settlement: red squirrel dispersal in contrasting habitats. - Journal of Animal Ecology 73: 1024–1034.

Hawkes, C. 2009. Linking movement behaviour, dispersal and population processes: is individual variation a key? - Journal of Animal Ecology 78: 894–906.

Heinz, S. K. and Strand, E. 2006. Adaptive Patch Searching Strategies in Fragmented Landscapes. - Evolutionary Ecology: 113–130.

Holyoak, M. et al. 2008. Trends and missing parts in the study of movement ecology. - PNAS 105: 19060–19065.

Jongejans, E. et al. 2008. Dispersal, demography and spatial population models for conservation and control management. - Perspectives in Plant Ecology, Evolution and Systematics 9: 153–170.

Kendall, B. E. and Fox, G. A. 2003. Unstructured Individual Variation and Demographic Stochasticity. - Conservation Biology 17: 1170–1172.

Knowlton, J. L. and Graham, C. H. 2010. Using behavioral landscape ecology to predict species' responses to land-use and climate change. - Biological Conservation 143: 1342–1354.

Kun, Á. and Scheuring, I. 2006. The Evolution of Density-Dependent Dispersal in a Noisy Spatial Population Model. - Oikos 115: 308–320.

Lambin, X. et al. 2012. High connectivity despite high fragmentation: iterated dispersal in a vertebrate metapopulation. - In: Dispersal Ecology and Evolution. Oxford University Press, in press.

Le Galliard, J.-F. et al. 2012. Patterns and processes of dispersal behaviour in arvicoline rodents. - Mol Ecol 21: 505–523.

Lima, S. L. and Zollner, P. A. 1996. Towards a behavioral ecology of ecological landscapes. - Trends in Ecology & Evolution 11: 131–135.

Matthysen, E. 2005. Density-dependent dispersal in birds and mammals. - Ecography 28: 403–416.

Merckx, T. et al. 2003. The evolution of movements and behaviour at boundaries in different landscapes: a common arena experiment with butterflies. - Proceedings of the Royal Society of London. Series B: Biological Sciences 270: 1815–1821.

Metz, J. a. J. and Gyllenberg, M. 2001. How should we define fitness in structured metapopulation models? Including an application to the calculation of evolutionarily stable dispersal strategies. - Proceedings of the Royal Society of London. Series B: Biological Sciences 268: 499–508.

Morales, J. 2002. Behavior at Habitat Boundaries Can Produce Leptokurtic Movement Distributions. - The American Naturalist 160: 531–538.

Morales, J. M. and Ellner, S. P. 2002. Scaling up Animal Movements in Heterogeneous Landscapes: The Importance of Behavior. - Ecology 83: 2240–2247.

Morales, J. M. et al. 2010. Building the bridge between animal movement and population dynamics. - Philosophical Transactions of the Royal Society B: Biological Sciences 365: 2289–2301.

Nathan, R. et al. 2008. A movement ecology paradigm for unifying organismal movement research. - PNAS 105: 19052–19059.

Neubert, M. G. and Caswell, H. C. 2000. Density-dependent vital rates and their population dynamic consequences. - J Math Biol 41: 103–121.

Olden, J. D. et al. 2004. Context-dependent perceptual ranges and their relevance to animal movements in landscapes. - Journal of Animal Ecology 73: 1190–1194.

Ovaskainen, O. 2004. Habitat-Specific Movement Parameters Estimated Using Mark–Recapture Data and a Diffusion Model. - Ecology 85: 242–257.

Ovaskainen, O. and Cornell, S. J. 2003. Biased Movement at a Boundary and Conditional Occupancy Times for Diffusion Processes. - Journal of Applied Probability 40: 557–580.

Ovaskainen, O. and Crone, E. E. 2009. Modeling animal movement with diffusion. - Spatial Ecology in press.

Ovaskainen, O. et al. 2008. An empirical test of a diffusion model: predicting clouded apollo movements in a novel environment. - Am Nat 171: 610–619.

Palmer, S. C. F. et al. 2011. Introducing a “stochastic movement simulator” for estimating habitat connectivity. - Methods in Ecology and Evolution 2: 258–268.

Patterson, T. A. et al. 2008. State–space models of individual animal movement. - Trends in Ecology & Evolution 23: 87–94.

Pe’er, G. and Kramer-Schadt, S. 2008. Incorporating the perceptual range of animals into connectivity models. - Ecological Modelling 213: 73–85.

Pe’er, G. et al. 2011. Breaking Functional Connectivity into Components: A Novel Approach Using an Individual-Based Model, and First Outcomes. - PLOS ONE 6: e22355.

Phillips, B. L. et al. 2010. Life-history evolution in range-shifting populations. - Ecology 91: 1617–1627.

Poethke, H. J. and Hovestadt, T. 2002. Evolution of density–and patch–size–dependent dispersal rates. - Proceedings of the Royal Society of London. Series B: Biological Sciences 269: 637–645.

Poethke, H. J. et al. 2011. The ability of individuals to assess population density influences the evolution of emigration propensity and dispersal distance. - J Theor Biol 282: 93–99.

Revilla, E. and Wiegand, T. 2008. Individual movement behavior, matrix heterogeneity, and the dynamics of spatially structured populations. - PNAS 105: 19120–19125.

Revilla, E. et al. 2004. Effects of Matrix Heterogeneity on Animal Dispersal: From Individual Behavior to Metapopulation-Level Parameters. - The American Naturalist 164: E130–E153.

Ricketts, T. H. 2001. The matrix matters: effective isolation in fragmented landscapes. - Am Nat 158: 87–99.

Robert, A. et al. 2003. Variation among Individuals, Demographic Stochasticity, and Extinction: Response to Kendall and Fox. - Conservation Biology 17: 1166–1169.

Ronce, O. 2007. How Does It Feel to Be Like a Rolling Stone? Ten Questions About Dispersal Evolution. - Annual Review of Ecology, Evolution, and Systematics 38: 231–253.

Ruxton, G. D. and Rohani, P. 1998. Fitness-dependent dispersal in metapopulations and its consequences for persistence and synchrony. - Journal of Animal Ecology: 530–539.

Schtickzelle, N. and Baguette, M. 2003. Behavioural responses to habitat patch boundaries restrict dispersal and generate emigration–patch area relationships in fragmented landscapes. - Journal of Animal Ecology 72: 533–545.

Schultz, C. B. and Crone, E. E. 2001. Edge-Mediated Dispersal Behavior in a Prairie Butterfly. -

Ecology 82: 1879–1892.
Schurr, F. M. 2012. How random is dispersal? From stochasticity to process in the description of seed movement. - In: Clobert, J. et al. (eds), Dispersal Ecology and Evolution. Oxford University Press, pp. 240–247.
Shreeve, T. G. and Dennis, R. L. H. 2011. Landscape scale conservation: resources, behaviour, the matrix and opportunities. - J Insect Conserv 15: 179–188.
Smouse, P. E. et al. 2010. Stochastic modelling of animal movement. - Philosophical Transactions of the Royal Society of London. Series B, Biological Sciences: 2201–2211.
Stamps, J. A. 1988. Conspecific Attraction and Aggregation in Territorial Species. - The American Naturalist 131: 329–347.
Stamps, J. 2001. Habitat selection by dispersers: integrating proximate and ultimate approaches. - In: Clobert, J. et al. (eds), Dispersal. Oxford University Press, pp. 230–242.
Stamps, J. A. and Blozis, S. A. 2006. Effects of natal experience on habitat selection when individuals make choices in groups: A multilevel analysis. - Animal Behaviour 71: 663–672.
Stamps, J. A. et al. 2005. Search Costs and Habitat Selection by Dispersers. - Ecology 86: 510–518.
Stamps, Judy A. et al. 2009. How Different Types of Natal Experience Affect Habitat Preference. - The American Naturalist 174: 623–630.
1992. Animal Dispersal: Small mammals as a model (NC Stenseth and WZ Lidicker, Eds.). - Springer Netherlands.
Stevens, V. M. and Coulon, A. 2012. Landscape effects on spatial dynamics: the natterjack toad as a case study. - In: Dispersal Ecology and Evolution. Oxford University Press, in press.
Stevens, V. M. et al. 2006. Quantifying functional connectivity: experimental assessment of boundary permeability for the natterjack toad (Bufo calamita). - Oecologia 150: 161–171.
Travis, J. M. J. et al. 1999. The evolution of density-dependent dispersal. - Proc Biol Sci 266: 1837.
Travis, J. M. J. et al. 2010. Towards a mechanistic understanding of dispersal evolution in plants: conservation implications. - Diversity and Distributions 16: 690–702.
Travis, J. M. J. et al. 2011. Improving prediction and management of range expansions by combining analytical and individual-based modelling approaches. - Methods in Ecology and Evolution 2: 477–488.
Travis, J. M. J. et al. 2012. Modelling dispersal: an eco-evolutionary framework incorporating emigration, movement, settlement behaviour and the multiple costs involved. - Methods in Ecology and Evolution 3: 628–641.
Travis, J. M. J. et al. 2014. Dispersal and species' responses to climate change. - Oikos 122: 1532–1540.
Vandermeer, J. and Carvajal, R. 2001. Metapopulation dynamics and the quality of the matrix. - Am Nat 158: 211–220.
Vercken, E. et al. 2012. The importance of a good neighborhood: dispersal decisions in juvenile common lizards are based on social environment. - Behavioral Ecology: 1059–1067.
Vuilleumier, S. and Metzger, R. 2006. Animal dispersal modelling: Handling landscape features and related animal choices. - Ecological Modelling 190: 159–170.
Vuilleumier, S. et al. 2006. Effects of Cognitive Abilities on Metapopulation Connectivity. - Oikos 113: 139–147.
Wiegand, T. et al. 2005. Effects of Habitat Loss and Fragmentation on Population Dynamics. - Conservation Biology 19: 108–121.
Zheng, C. et al. 2009. Modelling dispersal with diffusion and habitat selection: Analytical results

for highly fragmented landscapes. - Ecological Modelling 220: 1495–1505.
Zollner, P. and Lima, S. L. 1997. Landscape-level perceptual abilities in white-footed mice : perceptual range and the detection of forested habitat. - Oikos 80: 51–60.
Zollner, P. and Lima, S. L. 1999. Search Strategies for Landscape-Level Interpatch Movements. - Ecology 80: 1019–1030.
Zollner, P. A. and Lima, S. L. 2005. Behavioral tradeoffs when dispersing across a patchy landscape. - OIKOS 108: 219–230.
